# Terrestrial land cover shapes fish diversity in major subtropical rivers

**DOI:** 10.1101/2023.10.30.564688

**Authors:** Heng Zhang, Rosetta C. Blackman, Reinhard Furrer, Maslin Osathanunkul, Jeanine Brantschen, Cristina Di Muri, Lynsey R. Harper, Bernd Hänfling, Pascal A. Niklaus, Loïc Pellissier, Michael E. Schaepman, Shuo Zong, Florian Altermatt

**Affiliations:** Department of Evolutionary Biology and Environmental Studies, University of Zurich, Winterthurerstrasse 190, CH-8057 Zürich, Switzerland; Eawag, Swiss Federal Institute of Aquatic Science and Technology, Department of Aquatic Ecology, Überlandstrasse 133, CH-8600 Dübendorf, Switzerland; Department of Mathematics, University of Zurich, Winterthurerstrasse 190, CH-8057 Zürich, Switzerland; Institute of Computational Science, University of Zurich, Winterthurerstrasse 190, CH-8057 Zürich, Switzerland; Department of Biology, Faculty of Science, Chiang Mai University, Chiang Mai 50200, Thailand; National Research Council (CNR), Research Institute on Terrestrial Ecosystems (IRET), Lecce, Italy; The Freshwater Biological Association, The Hedley Wing, YMCA North Campus, Lakeside, Newby Bridge, Cumbria, LA12 8BD, UK.; Institute for Biodiversity and Freshwater Conservation, University of the Highlands and Islands, Inverness, UK.; Ecosystems and Landscape Evolution, Institute of Terrestrial Ecosystems, Department of Environmental Systems Science, ETH Zürich, Zürich, Switzerland; Unit of Land Change Science, Swiss Federal Research Institute for Forest, Snow and Landscape Research (WSL), Birmensdorf, Switzerland; Remote Sensing Laboratories, Department of Geography, University of Zurich, Winterthurerstrasse 190, CH-8057 Zurich, Switzerland

## Abstract

Freshwater biodiversity is critically affected by human modifications of terrestrial land use and land cover (LULC)^1,2^. Yet, knowledge of the spatial extent and magnitude of LULC-aquatic biodiversity linkages is still surprisingly limited, impeding the implementation of optimal management strategies^3^. Here, we compiled fish diversity data across a 160,000-km^2^ subtropical river catchment in Thailand characterized by exceptional biodiversity^4^ yet intense anthropogenic alterations^5^, and attributed fish species richness and community composition to contemporary terrestrial LULC across the catchment. We estimated a spatial range of LULC effects extending up to about 20 km upstream from sampling sites, and explained nearly 60 % of the variance in the observed species richness, associated with major LULC categories including croplands, forest, and urban areas. We find that integrating both spatial range and magnitudes of LULC effects is needed to accurately predict fish species richness. Further, projected LULC changes showcase future gains and losses of fish species richness across the river network and offer a scalable basis for riverine biodiversity conservation and land management, allowing for potential mitigation of biodiversity loss in highly diverse yet data-deficient tropical to sub-tropical riverine habitats.

## Introduction

Globally, human activities severely threaten biodiversity, challenging sustainable development goals proposed by the Intergovernmental Science-Policy Platform on Biodiversity and Ecosystem Services^6,7^. Biodiversity is declining, with losses of genetic, taxonomic, and functional diversity observed across all ecosystems, putting up to 1 million species at risk of extinction^6,8^. Freshwater ecosystems harbor an exceptionally high diversity of taxa^9^, supporting 43 % of all known fish species, with many drainage basins containing taxonomically and functionally unique fish assemblages^9,10^. Yet, many of these fish species are threatened worldwide, especially in riverine systems^11^. The decline of fish species richness has been widely attributed to major global change impacts in respective rivers, including modifications of river connectivity due to hydroelectric dams, warming and oxygen depletion of the water, overloading of nutrients, chemical pollution or direct exploitation by fishery^3,9^. Among all the factors, terrestrial land use and land cover (LULC) are recognized as a strong determinant for fish diversity and community distribution in most parts of the world^1,5^, with potential impact within a certain distance downstream. However, due to the limited spatial understanding of LULC-fish species associations, attributing current and predicting future effects of terrestrial LULC changes on fish diversity remains challenging, particularly in highly biodiverse yet data-deficient regions^12^. This impedes concrete and enforceable approaches to conservation management.

Riverine systems and surrounding terrestrial ecosystems are tightly interconnected at the catchment scale, resulting in cross-ecosystem linkages of resource and pollution flows^2^. LULC and its change thus impact river biodiversity through this terrestrial-aquatic coupling^13,14^. However, it remains difficult to estimate the spatial extent and magnitude of terrestrial LULC effects on riverine fish communities. For instance, croplands and urban areas, two typical LULC types associated with human activities, have strong effects on fish species richness in rivers, yet they are only reported at local scales or in qualitative manners^15-17^. Reported spatial ranges of the terrestrial LULC effects vary from dozens of meters to hundreds of kilometers downstream, yet the corresponding studies commonly focus on a few species and/or LULC types^16,18^.

Surprisingly, the fragmented mosaic structure and dynamic nature of LULC are regularly overlooked in assessments of terrestrial LULC impacts on highly biodiverse riverine systems. Combined, this leads to an inadequate understanding of terrestrial LULC effects on fish communities.

Large subtropical river catchments are global biodiversity hotspots, harboring among others a fascinating diversity of fish species^4^. One of these is the Chao Phraya River catchment in Thailand, holding many native yet threatened species such as the Siamese giant carp (*Catlocarpio siamensis*) or the endemic redtail sharkminnow (*Epalzeorhynchos bicolor*). This biodiversity, however, is threatened by anthropogenic changes, including the intensification and expansion of agricultural activities over past centuries. Croplands today occupy almost all the plains, and currently even expand into hilly and mountainous regions, therefore reduce natural ecosystems such as forest and shrubland. Urban areas have also expanded rapidly in recent decades, with a four-fold increase in area from 1992 to 2016^19^. This centuries-long and currently accelerating impact is predicted to intensify even further in the coming decades, leading to a loss of forest cover and threatening biodiversity in this region^20^.

Here, we implement a spatially explicit model that incorporates LULC maps with environmental DNA (eDNA)-based fish diversity assessments to evaluate terrestrial LULC effects on riverine fish species richness in the Chao Phraya catchment. Specifically, we provide a quantitative assessment of the spatial range of the effects of major terrestrial LULC types on fish species richness, and quantify these effects across the river catchment. Further, we project past and future fish diversity using historical and predicted future LULC data, identifying river habitats of fish species of conservation concern.

## Results

### A spatially explicit model

Fish communities along the major river channels in the Chao Phraya catchment were sampled using river water eDNA collected under base-flow conditions (Fig. 1). The detailed procedures are described in the Materials and Methods section and in^21^. In brief, we sampled water from 39 sites in 2016, which was then filtered, DNA extracted, and thereafter sequenced using two pairs of 12S primers (Kelly primers for vertebrates and MiFish primers for fish)^22,23^.

**Figure 1.**
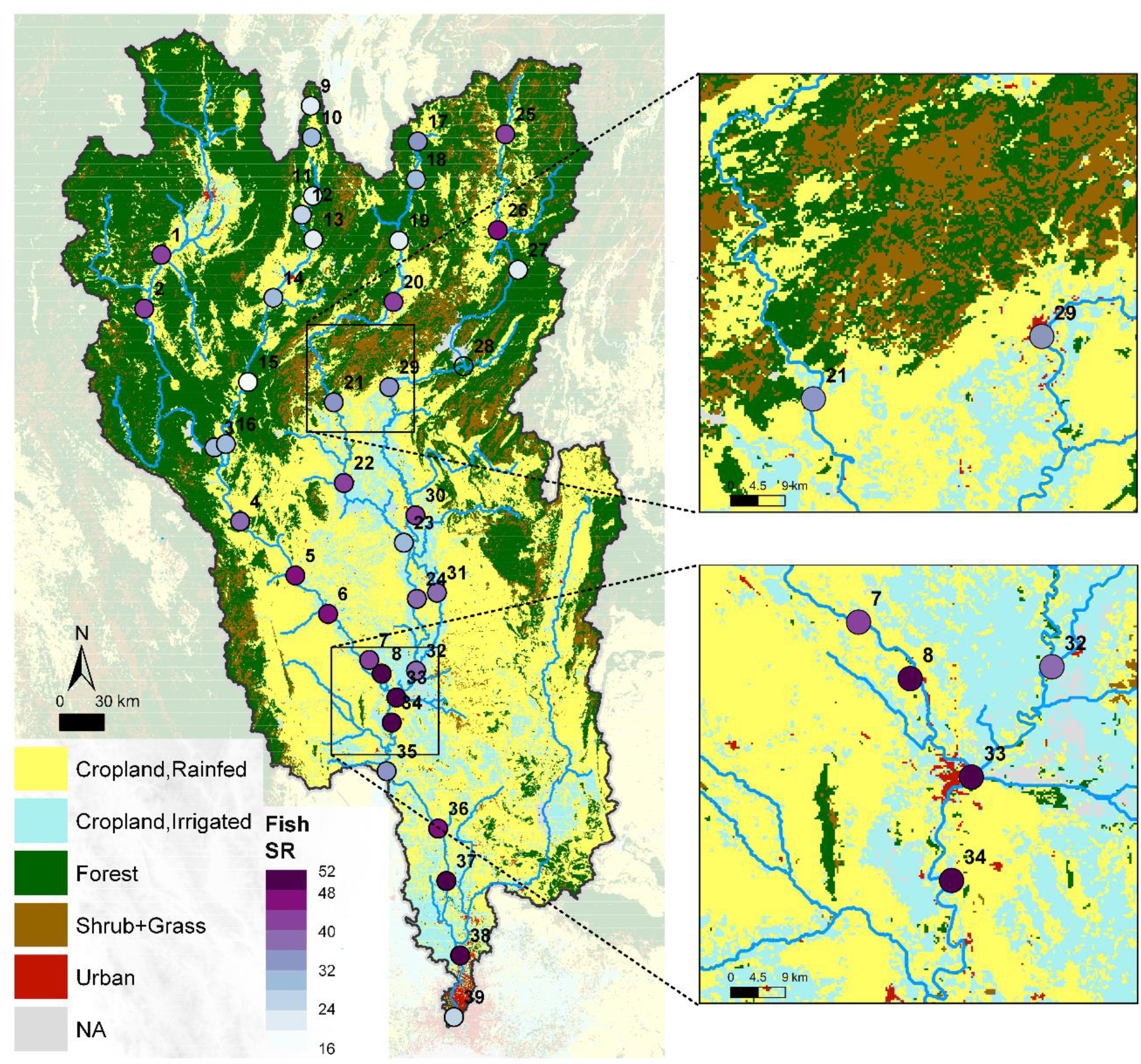
Riverine fish species richness (SR) and terrestrial land use and land cover (LULC) in the 160,000 km^2^ Chao Phraya catchment in Thailand. Fish species richness of 39 sites derived from environmental DNA in the major river channel is shown in purple dots, representatively covering the whole catchment. Zoom-in figures depict how the urban LULC type, with overall little area coverage, is especially predominant close to the major river channels.

From these two primer sets, we obtained in total 5,825,212 and 4,927,576 reads, respectively, which were merged by taking maximum read counts and matched with a total of 108 fish taxa (mostly at the species level, and subsequently referred to as fish species). At each site, species-level records were converted into presence/absence to calculate species richness. Among these fish species, seven were identified as critically endangered (CR), endangered (EN), vulnerable (VU), or near threatened (NT), according to the Red List of the International Union for Conservation of Nature (IUCN), after having removed alien and invasive species. Across the catchment, fish species richness ranged from 13 to 52 (34.5 on average), with strong variation across the river network (see also results reported in^21^). In general, fish species richness was high in the lower reaches of the Chao Phraya (Fig. 1), covering most of the plain area. In contrast, upstream reaches, generally in hilly or mountainous regions, showed low species richness but highly distinct communities among tributaries^21^. Nevertheless, sites in hilly or mountainous regions near croplands (site no. 1, 2, 4, 20, 25, and 26) showed relatively high species richness compared to adjacent sampling sites.

Terrestrial LULC was quantified using a 300 m-resolution land cover map (reference year 2016; European Space Agency Climate Change Initiative (ESA CCI)^19^). We reclassified the original 36 LULC classes into the five predominant and distinct LULC types: rainfed cropland (44 % cover), irrigated cropland (12 % cover), forest (36 % cover), shrub- and grassland (7 % cover), and urban (1 % cover) (Fig. 1; Table S1). Croplands are the dominant LULC type, covering 56 % of the catchment area, and are mainly found in the plains and along the rivers. Forest is predominantly found in the mountainous region at a higher elevation and includes a combination of broad-leaved evergreen and deciduous forests. Urban areas, though only occupying ∼1 % of the catchment area, are commonly found in the direct vicinity of the major river channels. They thus have a high potential to influence riverine fish species composition.

For each eDNA sampling site, we produced a map of flow distances based on the three-arc-second resolution HydroSHEDS (version 1) flow direction map^24^, i.e. we determined, for each pixel, the distance of the water flow path that connects it to the sampling site. These flow distance maps were then resampled to match with the ESA CCI land cover map. Overall, the flow distances to the sampling sites ranged from zero (for the pixel at the site itself) to 31— 1,076 km upstream (Fig. S1). In addition, we extracted river discharge, a proxy for fish diversity and river characteristics, from the HydroSHEDS database as a predictor of baseline fish species richness (Fig. S2)^25,26^.

Combining riverine fish diversity, LULC, and catchment data, we created a spatially-explicit model (hereafter referred to as the FishDiv-LULC model) by considering the spatial range and magnitude of effects from terrestrial LULC types on fish species richness. This model linked observed fish species richness to the terrestrial LULC effects integrated upstream along flow paths, and explained 58.7 % (adjusted R^2^) of the variance in the observed fish species richness (Table 1, see Materials and Methods). LULC-only and river discharge-only effects explained 21.7 % and 9.0 %, respectively, of the total variance in fish species richness, demonstrating significant terrestrial LULC effects on fish diversity in this catchment (Fig. S3).

**Table 1.** Estimations of FishDiv-LULC model parameters including terrestrial land use and land cover (LULC) effect values on riverine fish species richness and the spatial range *r* by flow distance. The magnitude of terrestrial LULC effect of rainfed cropland, irrigated cropland, forest, shrub- and grassland (S. + G.), and urban areas, respectively, are provided. *ℓ* is the log-likelihood function for optimization. The significance of parameters is determined by a likelihood-ratio test (see Materials and Methods and Supplementary Text). The uncertainty estimation is indicated in Table S7. We found a significant relative positive effect of rainfed cropland on fish species richness, but significant relative negative effects of forest and urban areas across the Chao Phraya catchment.

| | a | Crop (R) | Crop (I) | Forest | S. + G. | Urban | b | $r$ (km) | adj.$R^2$ | $-2\ell$ |
| --- | --- | --- | --- | --- | --- | --- | --- | --- | --- | --- |
| Value | 20.585 | 1.438 | -0.238 | -2.163 | -4.857 | -3.684 | 3.550 | 19 | 0.587 | 250.613 |
| p | — | 0.054 | 0.852 | 0.055 | 0.631 | 0.019 | 0.003 | <0.001 | — | — |

The estimated spatial range of terrestrial LULC effects on local fish species richness was 19 km (90 % CI: 11—34 km) upstream from sampling sites, suggesting a scale at which terrestrial LULC effects modulate fish species richness in the river. Rainfed cropland had a significant relative positive effect on fish species richness, whereas forest and urban areas had relatively negative effects (Table 1).

These relative positive or negative effects are the deviations to the baseline estimation across the catchment, and can be explained by differences in river nutrient availability among cropland and natural habitats^27,28^. To support this, we calculated river chlorophyll-a content (Chl-a), total suspended solids (TSS), and dissolved organic carbon (DOC), water properties reflecting nutrient availability and productivity, using Sentinel-2 level 2A data and found that rivers near croplands received stronger resource subsidies from surroundings and therefore supported larger fish communities and diversity (Fig. 2, see Materials and Methods). In addition, the centuries-long agricultural practices in these areas may have already pre-selected fish communities tolerant to terrestrial impacts from crop farming, such as high nutrient run-offs. Contrastingly, wastewater pollution and high anthropogenic disturbances in urban areas affect fish assemblage structure and can cause a decrease of fish species richness in rivers^29^.

**Figure 2.**
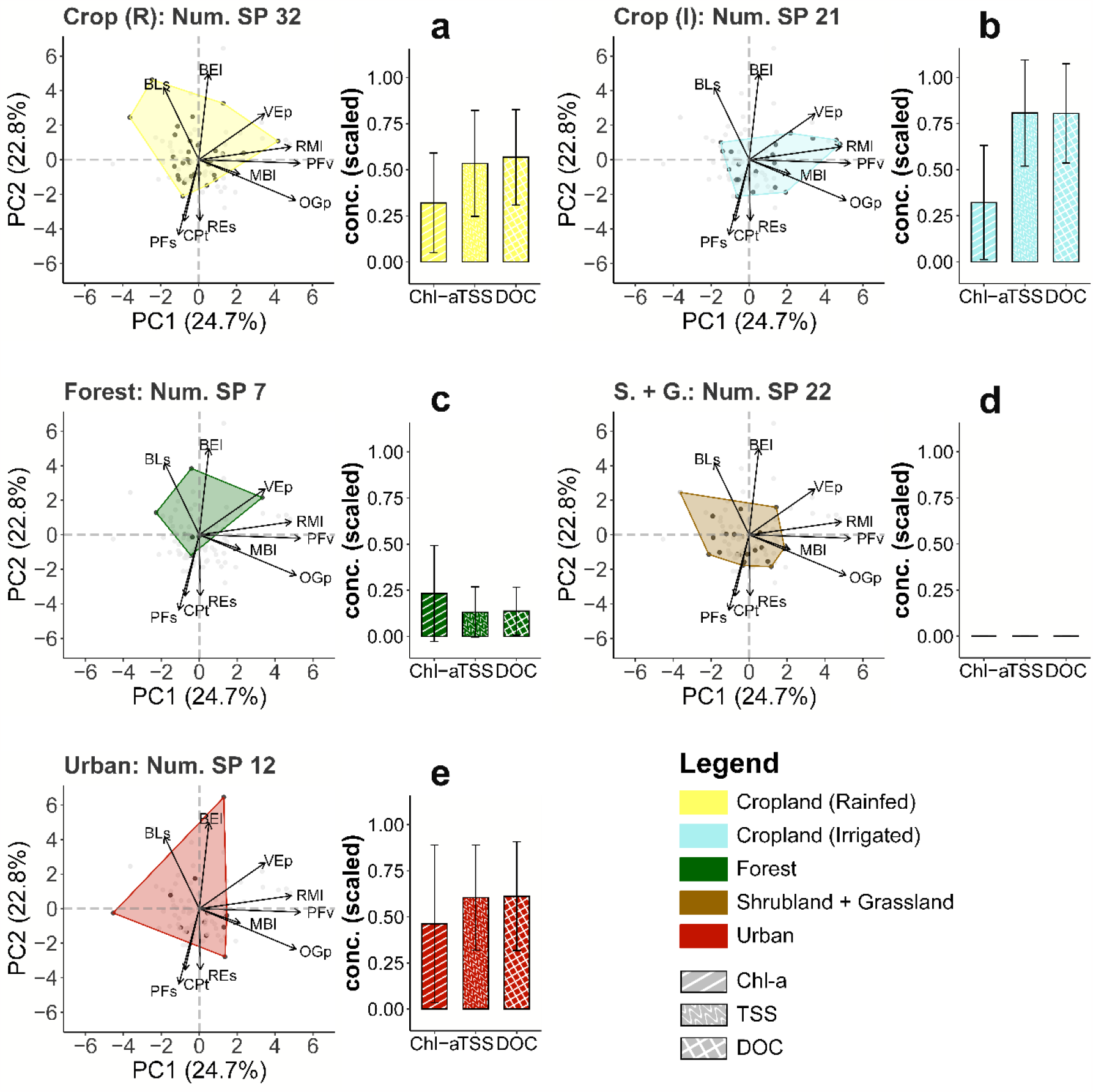
Fish species functional traits and water properties including chlorophyll-a (Chl-a), total suspended solids (TSS), and dissolved organic carbon (DOC) in relation to rainfed cropland (**a**), irrigated cropland (**b**), forest (**c**), shrub- and grassland (**d**), and urban (**e**) land use land cover (LULC) types, respectively, across the Chao Phraya catchment. The associated LULC for fish species was determined by the highest LULC effect value from species-level (presence/absence) modeling. The number of species (Num. SP) associated to each LULC type is shown in the figure. The trait space envelopes of LULC-associated fishes were created based on FISHMORPH database with a principal component analysis of functional trait space of fish species. Maximum body length (MBl), body elongation (BEl), vertical eye position (VEp), relative eye size (REs), oral gape position (OGp), relative maxillary length (RMl), body lateral shape (BLs), pectoral fin vertical position (PFv), pectoral fin size (PFs), and caudal peduncle throttling (CPt) were used to create fish trait space. Distinct fish trait space envelopes are observed among five LULC types. Water properties were estimated from Sentinel-2 data based on the major river channel pixels with a river width >60 m, and were min-max scaled for plotting. The error bars indicate the standard deviation. These figures illustrate that rivers in forested areas tend to have lower Chl-a, TSS, and DOC values, thereby less resource or nutrient subsidies compared with cropland and urban area.

### Robustness and mechanisms

We assessed the high robustness of our findings by comparing estimation results in two spatially separated sub-regions, splitting the fish sampling dataset in half. We separated 19 out of the 39 sites belonging to Northern Thailand in the hillier and more mountainous region with an elevation above 100 m; the remaining 20 sites were in Central Thailand, a plain-dominated region with an elevation below 100 m (Fig. S4, see Materials and Methods). For both datasets, we independently found similar positive terrestrial LULC effect from rainfed cropland, and negative effects from forest and urban areas, though the estimated spatial range of these effects differed between regions (Table S2). This indicated that the estimated LULC-fish species richness association was not driven by spatial clustering of the mountain and lowland region, but the LULC types themselves. It further indicated that even a smaller sampling effort (∼20 sites) was sufficient to effectively determine LULC effects. We further corroborated the observed terrestrial LULC effects by comparison to a null model. Specifically, we applied a neutral meta-community (NMC) model simulating fish species richness considering climate, fish habitat capacity, speciation, extinction, migration, and river network structure^26^, yet excluding any possible LULC effect. Our results showed that the FishDiv-LULC model (adjusted R^2^ = 0.587) explained a higher amount of variance than the NMC model (adjusted R^2^ = 0.255) and better captured fish species richness patterns in this subtropical region (Fig. S5 & Table S3), demonstrating strong terrestrial LULC effects on riverine fish species richness.

We then modified our FishDiv-LULC model into a species-level model, estimating terrestrial LULC effects on all individual fish species (see Materials and Methods). Next, we verified that terrestrial LULC directly drove fish species distributions through fish traits. To this end, we determined the associated terrestrial LULC type for each fish species through species-level modeling, then linked the determined LULC type to fish morphological traits extracted from the FISHMORPH database^30^, related to fish life cycle, ecology, and functional roles (see Materials and Methods). We analyzed envelopes of LULC type-associated fishes in trait space (in ordination space generated using a principal component analysis), and found distinct envelope shapes among different LULC types (Fig. 2). Fish species associated with rainfed cropland had high body elongation (BEl), high ranges of relative maxillary length (RMl), and body lateral shape (BLs), indicating better hydro-dynamics for swimming and a high trait diversity among these fishes; irrigated cropland-associated species showed high oral gape position (OGp) and RMl, relating to overall high trophic levels which coincided with high nutrient loadings from surroundings; forest-associated species had low maximum body length (MBl), relative eye size (REs), and caudal peduncle throttling (CPt), but high BLs, suggesting small body size but agility in swimming; urban-associated species had a high range of MBl and a large trait envelope area, indicating various strategies to adapt to high environmental and hydrological disturbances (Fig. 2, Table S4). Additionally, fish traits themselves were not significantly correlated with river network characteristics, such as river discharge, thus excluding a direct fish trait-river network linkage (Fig. S6). As a consequence, these results implied a strong linkage of terrestrial LULC on individual fish species distributions through fish traits and explained the formation of fish species richness patterns.

### Fish diversity projections with LULC changes

Conservation of riverine biodiversity is still short of effective methods to assess conservation potential of adjacent terrestrial land, and the latter’s quantitative impacts on riverine biodiversity. We show how the spatially explicit approach allows for projections of future richness and communities of riverine fish diversity, using minimal information accessible through global LULC products and river water eDNA sampling. To begin with, we produced a map of terrestrial LULC effects on riverine fish species richness, which can be understood as the accumulative fish species richness in the river due to LULC effects from the terrestrial land (Fig. 3a, see Materials and Methods). We then projected fish species richness in the major river channels, where regions rich in fish species were mostly observed in the plains and some hilly and mountainous regions close to the croplands, successfully capturing the spatial variation of fish species richness (Fig. 3b).

**Figure 3.**
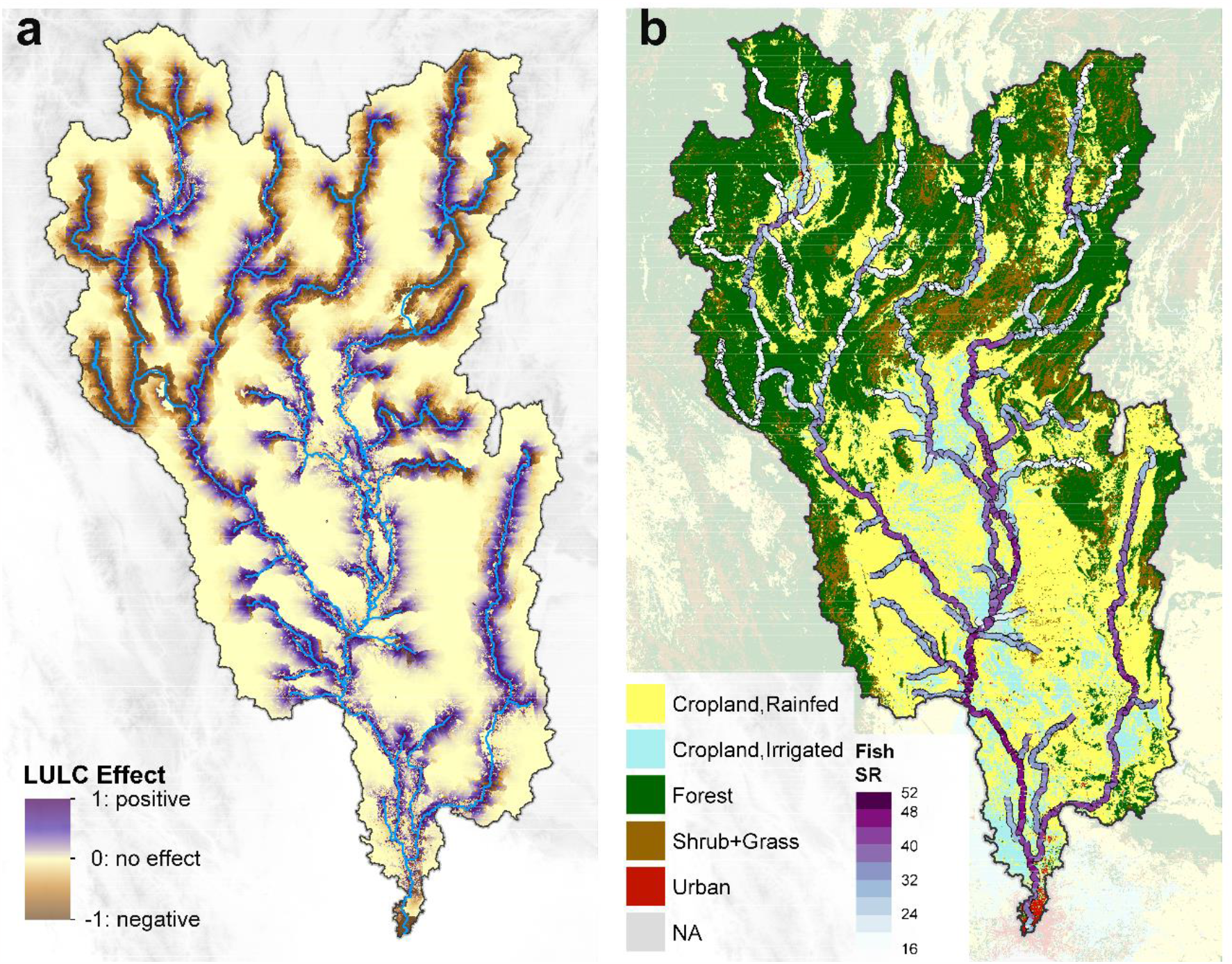
(**a**) Map of terrestrial LULC effects on riverine fish species richness in the main river channel of the Chao Phraya catchment. The effect value is the accumulative fish species richness change due to LULC impacts along the major river channels (unit: number of species × km) with a spatial resolution of 1 km^2^. (**b**) Projected pattern of riverine fish diversity (species richness, SR) in the major river channels of the Chao Phraya catchment. The LULC map is embedded as the background layer. The projection shows a high consistency with eDNA-derived fish diversity sampling in Fig. 1.

Anthropogenic LULC changes continue to intensify worldwide^31^. To explore potential impacts of such LULC changes, we used past and modeled future LULC maps to retrospect and forecast riverine fish species richness patterns. To isolate terrestrial LULC effects, we assumed that climate, flow discharge, and river network remained constant. For the past, 24 years of historically observed LULC change (1992—2016) was evaluated by adopting the ESA CCI land cover map of 1992^19^ (Fig. 4a); for the future, 34 years of modeled LULC changes under future scenarios (2016—2050) were assessed according to the products of GLOBIO4 scenario data^32^. Land use maps of 2050 under shared socio-economic pathway 1 and representative concentration pathway 2.6 (SSP1 RCP2.6), SSP3 RCP6.0, and SSP5 RCP8.5 scenarios were used for forecasting future fish diversity (Fig. 4d for SSP5 RCP8.5, Fig. S7 for other scenarios).

**Figure 4.**
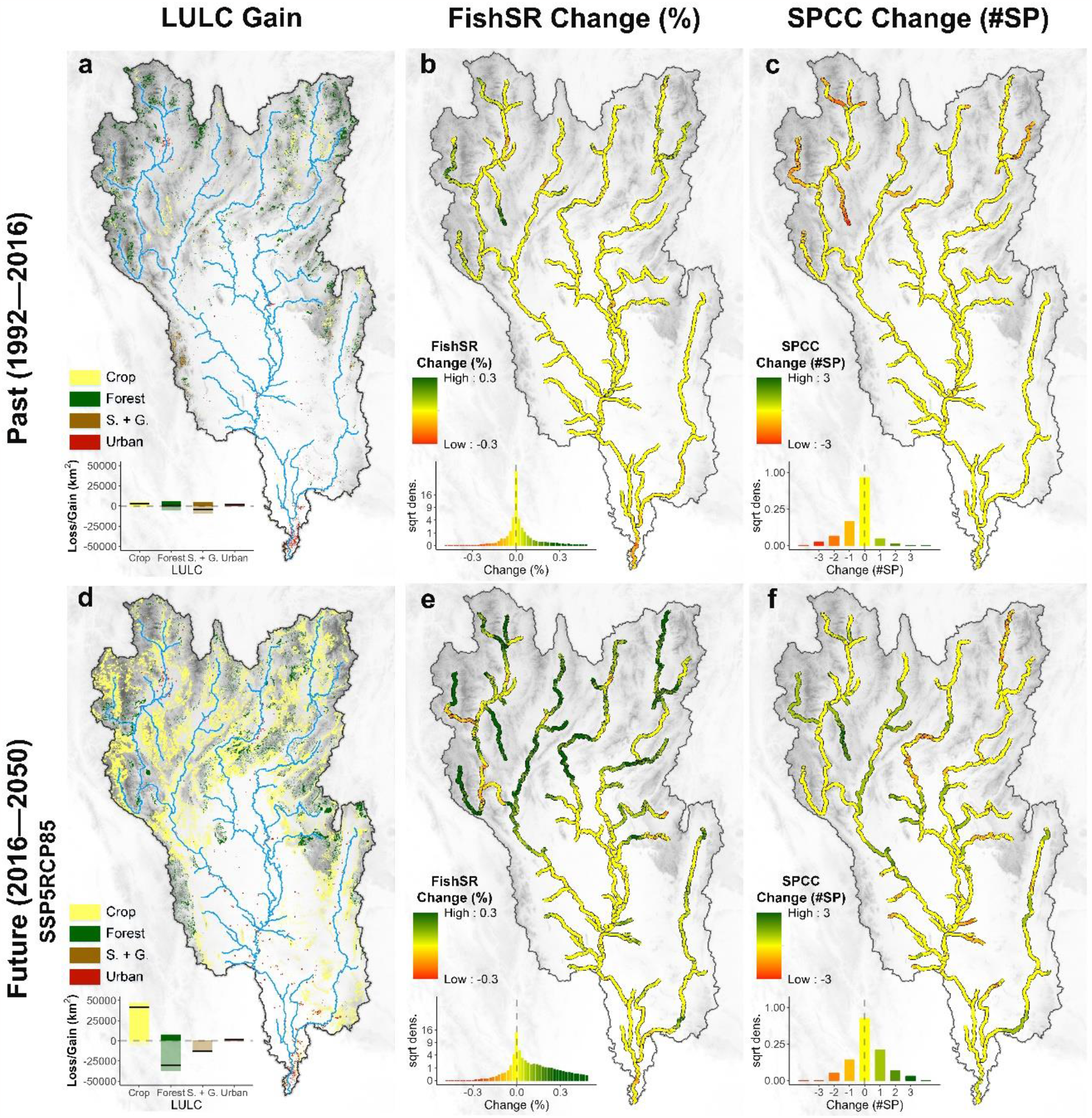
Predicted diversity changes of overall fish species richness (SR) and richness of fish species of conservation concern (SPCC) due to LULC changes in the periods of 1992—2016 (past) and 2016—2050 (future). Future prediction of LULC was adopted from GLIOBIO4 data, and the list of seven SPCC was derived from the Red List of the International Union for Conservation of Nature (IUCN) with alien or invasive fish species removed. We observed croplands and urban areas expansion over the past (**a**) and future (**d**) periods, resulting in a reduction of natural habitats such as forest, shrub- and grassland, especially in the mountainous region in the period of 2016—2050. Projected percentages of overall fish species change between 1992 and 2016 (**b**), and between 2016 and 2050 (**e**). Projected numbers of SPCC change between 1992 and 2016 (**c**), and between 2016 and 2050 (**f**). These predictions demonstrate different trends of fish species richness change between overall fish species richness and SPCC richness. Please refer to Fig. S6 for other scenarios SSP1RCP26, SSP3RCP60.

According to these LULC maps, urban areas increased by 295 % from 1992 to 2016, and were forecasted to increase by another 73 % from 2016 to 2050 (under SSP5 RCP8.5 scenario), mostly along major river channels. Moreover, predicted LULC changes in the future contained a conversion of forest, shrub- and grassland to croplands, leading to a 25 % increase in croplands and a 26 % decrease (under SSP5 RCP8.5 scenario) in forest, mostly in the hilly and mountainous region. Based on these scenarios, we predicted fish species richness for 1992 (past) and for 2050 (future; under the three scenarios) (see Materials and Methods), and produced maps of percentage of fish species richness changes (Fig. 4b for the past, Fig. 4e for the future under SSP5 RCP8.5, Fig. S7 for other scenarios). From 1992 to 2016, we observed only slight changes in fish species richness (Fig. 4b), partly due to local urbanization and LULC change from shrub- and grassland to forest (a 21 % decrease in shrub- and grassland); yet, from 2016 to 2050, we forecast a remarkable increase of fish species richness in the hilly and mountainous region under three scenarios (7.9—14.3 %) because of a strong expansion of croplands by 29—54 % in that region (Fig. 4e under SSP5 RCP8.5, Fig. S7 for other scenarios). However, these predictions about species richness did not reflect impacts of terrestrial LULC on less-common and endangered fish species, but mostly on already benefited common or generalist fish species.

We therefore repeated our projections, this time using the species-level model and focusing on the subset of seven fish species that were of conservation concern (SPCC) (Table S4, Fig. 4c & f under SSP5 RCP8.5, Fig. S7 for other scenarios; see Materials and Methods). Not unexpected, we found that the patterns of change in SPCC richness diverged from the patterns of change in overall fish species richness, with a high inconsistency in the hilly and mountainous region. This indicated that river channels with the most intense LULC change from natural habitats to croplands tended to lose SPCCs, emphasizing the necessity of land management regulations to mitigate such effect.

## Discussion

We demonstrate how a spatially explicit model integrating terrestrial LULC and eDNA-based fish diversity allows for attributing and forecasting of terrestrial LULC effects on riverine fish species richness in a large subtropical catchment. Specifically, for overall fish species richness in the Chao Phraya catchment in Thailand, we found a relative positive terrestrial LULC effect from rainfed cropland, most likely caused by high resource and nutrients subsidies, but relative negative effects of forest and urban areas, with a maximal distance of up to 19 km upstream from sampling sites. By analyzing fish traits in relation to specific terrestrial LULC types, we derived characteristic LULC-fish trait linkages and thereby explained the possible formation of fish species richness patterns. Furthermore, forecasts over future LULC change scenarios indicated strong human impacts on future fish diversity patterns, especially in the hilly and mountainous region where cropland expansion would increase fish species richness. In contrast, fish species of conservation concern would be negatively influenced by such LULC changes. Our approach can be applied to other biomes worldwide and allows for attributing fish diversity and its changes to major anthropogenic LULC changes.

The relative positive effect of rainfed cropland on fish species richness that we found align with previous case studies which reported increased fish species richness because of agricultural activities and associated nutrient subsidies, for instance, in Northern Europe and Southern Brazil^15,33^. To show how differences in river nutrient availability influence fish species richness in relation to LULC types, we calculated proxies for river nutrient and productivity, namely Chl-a, TSS, and DOC, for major river channel pixels (see Materials and Methods). In this computation, river channels narrower than 60 m were excluded because no RS in-river values were available. We found higher Chl-a, TSS, and DOC values close to croplands compared to forests (Fig. 2). These data provide evidence that rivers near croplands received high nutrient run-offs from the surrounding terrestrial land, which subsequently resulted in increases in algal biomass and food availability. With enlarged resource availability, especially for more generalist and omni- and algivorous fish species, rivers near cropland can consequently harbor more fish species^15^.

Importantly, the observed relative positive terrestrial LULC effects from rainfed cropland represented an averaged deviation to the baseline species richness estimation across the whole catchment. This baseline estimation, as suggested, included the effect of river characteristics (e.g., river discharge), the species pool, and an averaged terrestrial LULC effect on fish species richness across the catchment. As such, it did not suggest that cropland expansion would always benefit fish species richness, and would also promote highly specialist fish species which are often of conservation concern. For instance, we observed a hump-shaped relationship between fish species richness and river Chl-a, TSS, and DOC proxies indicating higher nutrient availability from agriculture (Fig. 5). When differentiating the mountainous and plain sites, the mountainous sites were always on the increasing slope of the peak, meaning that species richness may still increase with further cropland expansion. Conversely, fish species richness started to decline in the plains where nutrient loadings and intensity of agriculture were already very high. Furthermore, we found no significant positive effects of irrigated cropland, which was a more intense form of agriculture and often associated with excessive nutrient loadings and heavy use of pesticides and fertilizers^34^, likely with negative effects on fish diversity^35^. Ultimately, our analysis demonstrated a variety of cropland effects to fish species richness, implying the necessity of adequate and sustainable land management and agriculture regulations in this region.

**Figure 5.**
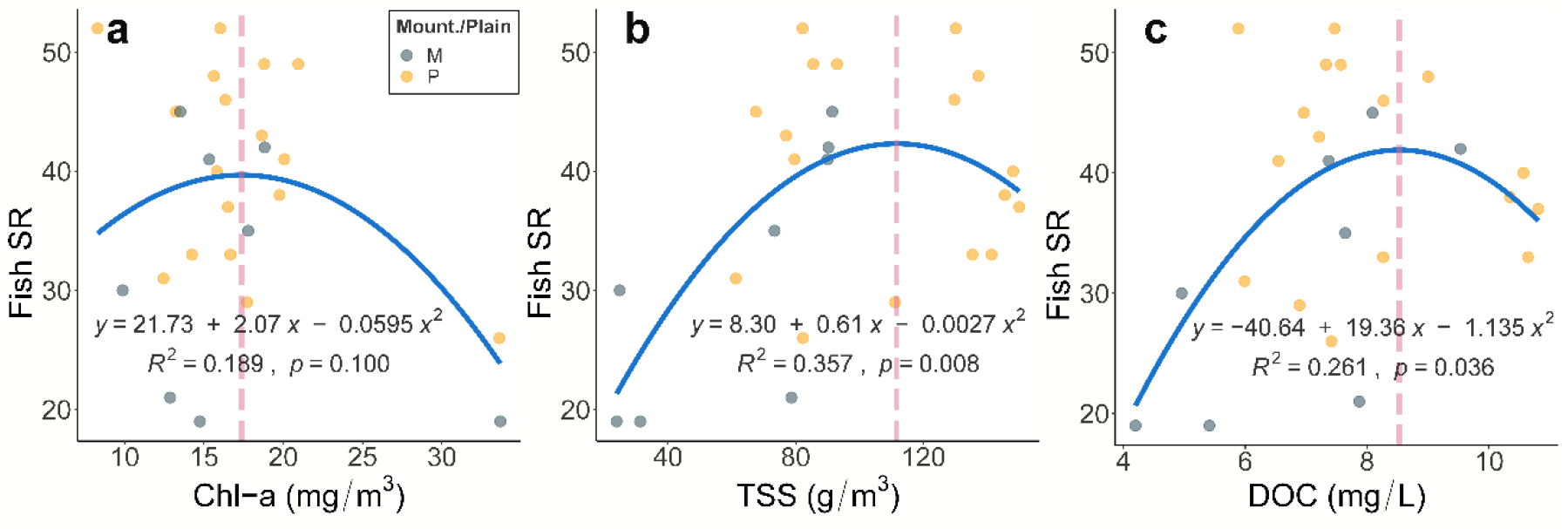
Relationship between eDNA-derived fish species richness and remote sensing (RS)-derived water properties of (**a**) chlorophyll-a (Chl-a), (**b**) total suspended solids (TSS), and (**c**) dissolved organic carbon (DOC) in the Chao Phraya catchment. Sampling sites in very narrow river channels were removed to ensure accuracy in RS calculations. Sites in the mountainous and plain areas are labeled in slate grey and gold, respectively. The blue line gives a quadratic fit to the data. The dashed line indicates maximum fish species richness (SR).

The expansion of cropland may also cause homogenization among riverine fish assemblages in the catchment. In the previous results, we focused on patterns of fish species richness, arguably the most commonly used metric of biodiversity^26^. However, this metric is insensitive to uniqueness within communities and has intrinsic limitations. Our further predictions over the future LULC change scenarios showed a decrease in the uniqueness of fish species across the catchment. When calculating the Jaccard similarity index of predicted fish assemblages between sites in the mountainous river and sites in the plain river from 2016 to 2050 under SSP5 RCP8.5 scenario, we found a remarkable increase in similarity of fish assemblages in the future, suggesting homogenization among fish assemblages in this catchment (Fig. S8, see Material and Methods). In general, natural habitats have good conservation potential for specialist and narrow-ranged fish species. For example, river channels near forests can harbor more rare fish species, essentially contributing to local biodiversity^36^. Nevertheless, due to cropland expansion into the hilly and mountainous region, native and often endangered species associated with natural habitats could be replaced by cropland-associated and/or wide-ranging species, such as *Osphronemus goramy* or *Boesemania microlepis*, causing a loss of uniqueness and homogenization^33^. Consequently, our future predictions do not suggest that fish diversity loss could be mitigated in the future, but imply that specific land management and regulations are needed to alleviate adverse impacts of LULC on less-common and endangered species.

Our approach has strong potential to be applied to any river catchment, given high cost-efficiency as well as comparability of results of river eDNA sampling and globally available high-resolution LULC products. In our model, the baseline estimation of fish species richness can be changed to other factors determining fish species richness patterns, such as temperature, habitat size or drainage area, geological and/or historical events, river water properties, and river structures^25,26^. In the Chao Phraya catchment, we considered flow discharge as a proxy for water balance, stream order, and fish habitat capacity. We did not include temperature as a parameter, because it was mostly homogenous in this region (Fig. S2a). However, other long rivers, such as the Mississippi, Yangtze, or Danube, flow through multiple biomes and have a larger elevation range; therefore, for those, it may be necessary to include climatic factors and/or river water properties in the baseline fish species richness estimation.

Human modifications of terrestrial landscapes are a primary driver of LULC change, and through cross-ecosystem linkages, are continually reshaping riverine biodiversity. Cropland and urban area, the typical anthropogenic LULC types having pronounced effects on this biodiversity, can be directly governed by legislation and policies through controlling the area and position in the landscape. Therefore, when developing adequate conservation strategies for freshwater ecosystems, we need careful considerations of current and future LULC distributions. In this sense, global initiatives, such as the 30 by 30 Initiative^37^, aiming to manage the specific use of land, should not only consider the total amount of land but also its spatial position and the underlying cross-ecosystem effects at the catchment level. Our approach provides precise estimation of local fish diversity changes under anthropogenic terrestrial LULC alterations, giving both scientists and stakeholders a potent tool in land management and conservation area design.

## Methods

The study was conducted in the Chao Phraya River catchment located in Northern and Central Thailand, covering rivers in both mountainous and plain landscapes. We combined fish diversity data from eDNA sampling in the rivers (elevation ranging from 2 m to 509 m a.s.l.) and a land use and land cover (LULC) map across the 160,000 km^2^ catchment (Fig. 1).

### Environmental DNA sampling and fish data

The fish data were derived from an eDNA metabarcoding study with methodological details published therein^21^. Briefly, eDNA sampling was carried out in 2016 at 39 sites in major river channels, during the dry season under base-flow conditions. At each site six samples were collected from the left bank, channel center, and right bank, respectively (two replicates each; 234 samples in total). Following on-site filtration (600 mL water in total per sampling site), DNA was extracted using standardized methods, and metabarcoding analyses were carried out using two separate molecular assays based on the mitochondrial 12S region, subsequently referred to as the Kelly primers for vertebrates and the MiFish primers for fish^22,23^. To improve accuracy of sequence assignment, we created a customized reference library database from GenBank targeting fish species known to occur in the Chao Phraya River catchment, according to OEPP Biodiversity Series Vol. 4 Fishes in Thailand^38^ and the Checklist of Freshwater Fishes of Thailand (http://www.siamensis.org/). During bioinformatic processing, sequences were assigned to a total of 108 fish taxa (mostly at species level, so referred to as fish species), with 82 and 93 taxa recovered using the Kelly primers and MiFish primers, respectively. Differences in species communities between the two assays can largely be accounted for by unequal representation of the respective DNA regions in the reference database and differences in species level resolution^21^. We merged these two data sets by choosing the higher read counts at each site for each fish species and calculated species richness. The list considered included native and naturalized species, part of which were also used in aquaculture. We further matched the detected fish species with the Red List of the International Union for Conservation of Nature (IUCN), having removed any alien or invasive species. In total, seven species were identified as critically endangered (CR), endangered (EN), vulnerable (VU), or near threatened (NT), and were treated as species of conservation concern (SPCC) in our fish data (Table S5).

### Land use and land cover (LULC) data

We used the European Space Agency Climate Change Initiative (ESA CCI) land cover map, with an yearly interval (website: https://www.esa-landcover-cci.org/)^19^. Specifically, we used the 300 m resolution map from 2016 to temporally match our fish data. Among the 36 classes in the classification system, 22 classes—including croplands, forests, shrublands, grasslands, and urban areas—were observed in the Chao Phraya catchment.

To alleviate uncertainties from rarely observed LULC types, we excluded or merged those LULC types occupying <0.2 % of the area. Further, we removed all areas covered by water (lakes, reservoirs, and rivers), such that only terrestrial LULC types were used. We then merged the LULC map to a five-class system, comprised of rainfed cropland, irrigated cropland, forest, shrub- and grassland, and urban areas (Fig. 1). A detailed recoding table is shown in Table S1.

### River channel and catchment data

To improve spatial modeling performance, we adopted the three-arc-second resolution (∼92 m at the equator) HydroSHEDS (version 1) flow direction map to calculate potential catchments for sampling sites^24^. A pixel in a flow direction map contains one of eight flow directions indicating an adjacent pixel to where water flows. Therefore, for each sampling site, we produced a catchment map in which we tracked along the flow direction map to find upstream pixels and calculated the flow distance with the haversine formula (see Supplementary Text). The catchment maps with flow distance were resampled to match with LULC data and then were used in the FishDiv-LULC model introduced below as *d*_ij_ (the flow distance between a catchment pixel *j* and sampling site *i*). The major river channels (blue lines in Fig. 1) were extracted using a threshold drainage area of ∼810 km^2^ (100,000 pixels). We also extracted the river discharge (*Q*) for all sampling sites and major river channel pixels from the HydroSHEDS database.

### Modeling terrestrial LULC effects on fish species richness (FishDiv-LULC model)

We developed a spatially explicit modeling framework to assess terrestrial LULC effects on riverine fish species richness, considering both the spatial range and magnitude of LULC effects. Let *k* = 1, 2, …, *K* represent the LULC type, and *j* = 1, 2, …, *N*_ik_ represent the pixel index of LULC type *k* in the catchment map of sampling site *i* (*i* = 1, 2, …, *M*; *M* = 39 in this case).

Specifically, we assume that the fish species richness at sampling site *i* (*B*_i_) equals to the sum of effects of different LULC types in its catchment (∑ S_ik_) plus a baseline prediction *B*_i_^0^ and an error *ε*_i_ (Eq. 1).

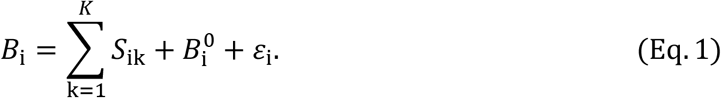

Then, for each site of interest (e.g., site *i*), the effect of a pixel *j* with LULC type *k* on the fish species richness can be written as *V*_k_ · *A*_j_ · *f*(*d*_ij_). Whereby *V*_k_ is the magnitude of the effect of LULC type *k, f*(*d*) is a distance decay function, and *A*_j_ is the area of pixel *j* depending on the coordinates and is estimated by the haversine formula (see Supplementary Text).

We evaluated five commonly-used distance decay functions, with the widely-used exponential decay function performing the best (see Supplementary Text and Table S6). In the exponential distance decay function, distance is the flow distance in the catchment map (*d*_ij_) from pixel *j* to site *i* (Eq. 2).

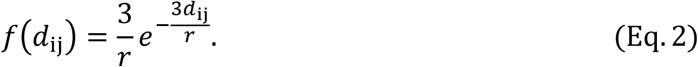

Here, the parameter *r* indicates the effective distance at which the magnitude of terrestrial LULC effect has dropped to ∼5% of its original value. Consequently, the terrestrial LULC effect is explicitly expressed in Eq. 3.

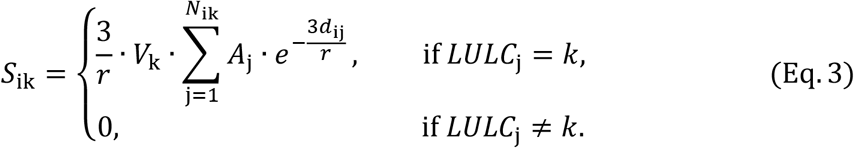

Note that 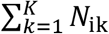 is equal to the total number of pixels in the catchment map of site *I* (i.e., every pixel belongs to one specific LULC type *k*). The base-line prediction *B*^0^ is expressed using river discharge (*Q*), which best explains fish species richness pattern (Fig. S2, Eq. 4).

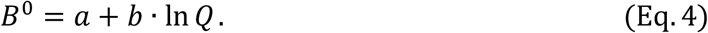

### Optimization of model parameters

Model parameters were estimated by solving a maximum likelihood problem, given the observed fish species richness *B*_i_. Subsequently, we write the above optimization problem explicitly as Eq. 5 (with vector-matrix notation).

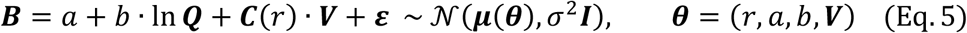

Whereby ***C***(*r*) is an *M*-by-*K* matrix with elements 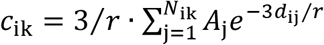 depending on the distance parameter *r*; ***V*** is a *K*-vector of magnitude parameters *V*_k_; ***B*** is an *M*-vector of observed fish species richness *B*_i_; ***ε*** is an *M*-vector of errors, in which each element is assumed independent and identically normally distributed; *a* and *b* are constants, with *a* being the intercept in the estimation; ***Q*** is an *M*-vector of river discharge values; and **θ** is a *K*+3 dimensional parameter.

Then, we estimate **θ** by maximum likelihood. We explicitly write the log likelihood function (*ℓ*) as Eqs. 6—7.

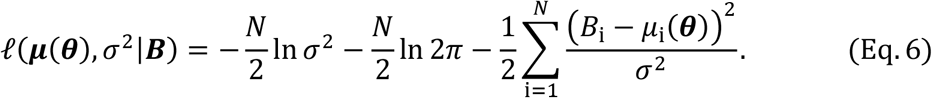

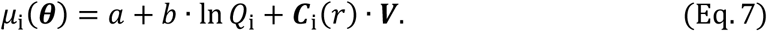

Whereby, ***C***_i_(*r*) is the *i*^th^ row of matrix ***C***(*r*), and *σ*^2^ is the variance to estimate. The optimal parameter vector 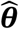 is determined by maximizing *ℓ*, or equivalently, minimizing −2*ℓ*. We also computed adjusted R^2^.

### Correlation and significance of model parameters

Variance inflation factor (VIF) and paired Pearson’s correlation of parameters were calculated under the estimated effective spatial range (*r* = 19 km). The correlation among the estimated terrestrial LULC effects is relatively weak (Table S7 and Fig. S9). The significance of these effects was determined by a likelihood-ratio test (see Supplementary Text).

### Validation, robustness, and estimation of uncertainties

We used leave-one-out cross-validation to assess the uncertainty of our parameter estimates. Specifically, we reserved one sampling site from the fish data for testing, followed by estimating the parameters based on the remaining 38 sites (i.e., the training set). Then, we predicted the fish species richness value on the reserved site and compared the predicted value with real observation. We repeated the whole process for each site and calculated the root-mean-square error (*RMSE*) of our model to be 8.02 (mean of prediction: 34.9).

The robustness of our model was tested by splitting the sampling sites into mountainous sites (elevation > 100 m, 19 sites), and the remaining sites in the plains (20 sites). Then, we fitted the same model to the two data subsets (Table S2).

We also plotted the residuals of the model against terrestrial LULC effects and the model prediction (Fig. S10), and we did not find any obvious trend in these scatter plots. To estimate the uncertainty of parameters, we calculated profile likelihood-ratio confidence intervals (CI) of levels of 50 % and 90 % for each model parameter (see Supplementary Text; Table S8).

### Modeling terrestrial LULC effects on fish species distributions (species-level modeling)

To predict the habitat distribution of fish species, we generalized our model to a species distribution model by modifying the fish species richness observation of Eq. 5 with a logit function. Specifically, we substitute ***B*** with *ln*(***P***/1-***P***), where ***P*** is the probability of presence of a fish species (Eq. 8).

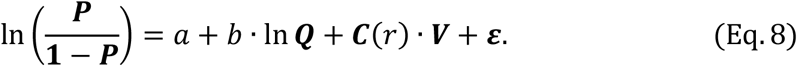

Then, we applied a maximum likelihood estimation to find the effective spatial range *r* and magnitude ***V*** of terrestrial LULC effects. The associated LULC for each species was assigned by the LULC type with the highest LULC effect value.

### Terrestrial LULC effect map

We mapped the LULC effect on fish species richness for each terrestrial pixel (*E*_LULCj_) by tracing the pixel location with the flow direction map and summing up its LULC effect (*S*_LULCj_) along the major river channels downstream, using the optimal *r* and ***V*** (Eq. 4). The integral with flow path (*L*_j_) starts from where pixel tracing entering the major river channels till the spatial range (*r*) downstream. Additionally, we scaled the map by dividing by the pixel area, so that the value in the map can be directly perceived as the accumulative change of fish species richness (unit: num. species × km) due to the terrestrial LULC effect of pixel location with a 1 km^2^ resolution (Eq. 9; see Fig. 3a).

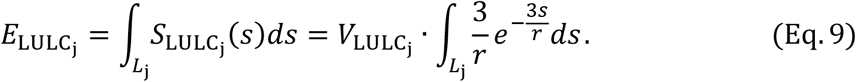

Based on the previous bootstrapped samples, we predicted the terrestrial LULC effect map 2,000 times and then calculated the interquartile range (IQR) as a metric of uncertainty (Fig. S11a & c).

### Neutral meta-community (NMC) model as a null model

We compared our result with a null model of a quasi-neutral river meta-community model, which considers climate, fish habitat capacity, speciation, extinction, migration, and river network structure^26^. This model uses meta-community theories and fish dispersal in the riverine network to predict fish species richness pattern. In the NMC simulation, the product of average annual runoff production (AARP) and watershed area (WA) was replaced by river discharge (Q) acquired from the HydroSHEDS data, as they represent similar meaning and have high correlations. We derived the optimal parameters for NMC model (Tab. S7) and the prediction error pattern (Fig. S3).

### Projecting riverine fish species richness map

We applied our model to major river channel pixels in the catchment to generate a riverine fish species richness map (Fig. 3b). To do so, we produced a local catchment within a 19 km spatial range for each major river channel pixel, followed by applying our FishDiv-LULC model to predict fish species richness in the river. We also assessed the uncertainty by predicting fish species richness based on the optimal solutions from 2,000 bootstrapped samples and then computed IQR of prediction results for major river channel pixels (Fig. S11b & d).

### Fish traits in relation to terrestrial LULC

We collected ten major fish morphological traits from the FISHMORPH database^30^. They are maximum body length (MBl), body elongation (BEl), vertical eye position (VEp), relative eye size (REs), oral gape position (OGp), relative maxillary length (RMl), body lateral shape (BLs), pectoral fin vertical position (PFv), pectoral fin size (PFs), and caudal peduncle throttling (CPt), relating to fish metabolism, hydro-dynamism, body size and shape, trophic levels and impacts, etc. Then, we linked fish traits and associated LULC type for each species according to the largest positive LULC effect value in the species-level modeling result. Species without valid species-level models were removed, so 93 out of 108 species were finally analyzed. Lastly, we mapped trait envelopes of LULC-associated fish species using a principal component analysis of ten morphological trait space.

### Fish species richness changes in the past and future

To predict fish species richness patterns in the past and future, we used ESA CCI land cover map in 1992 (the first year of the product) and GLOBIO4 land use maps in 2050 as past and modeled future LULC maps, respectively^19,32^. The GLOBIO4 2050 land use maps are predicted based on the present ESA CCI land cover map, showing good consistency with the global LULC product used in our modeling. For 2050, we used three LULC maps under shared socio-economic pathway 1 representative concentration pathway 2.6 (SSP1 RCP2.6), SSP3 RCP6.0, and SSP5 RCP8.5 scenarios.

Due to the lack of differentiation between rainfed cropland and irrigated cropland in the future maps, we merged these two LULC types (four LULC types in total) and re-fitted the model to predict fish diversity changes in the past and future. The new validation is shown in Table S9. We predicted riverine fish species richness maps of 1992 and 2050 (under three scenarios), and then calculated the percentage of changes in the periods of 1992—2016 (past) and 2016—2050 (future) (Fig. 4 & S6).

### Distribution changes of fish species of conservation concern (SPCC) in the past and future

We predicted the distribution map for each SPCC by firstly calculating a probability map in major river channels, and afterwards, determining presence/absence at each river channel pixel by a threshold with the highest true skill statistic value.

To assess the effects of LULC changes on SPCC, we also predicted the distribution maps for SPCCs in the past (1992) and forecast future fish distribution changes under three LULC change scenarios (2050). As a result, the distribution changes of SPCCs from 1992 to 2016 (past) and under three scenarios from 2016 to 2050 (future) are depicted in Figures 4 & S7.

### River water properties from remote sensing data

We estimated river water properties of chlorophyll-a content (Chl-a), total suspended solids (TSS), and dissolved organic carbon (DOC) to show nutrient/resource availability in rivers. To improve accuracy, image collections of Sentinel-2 level 2A surface reflectance (SR) data were used to obtain a 20-m cloud-free image on the Google Earth Engine. Then, we extracted major river channel pixels using a water occurrence map from the 30-m global surface water data with a threshold of 0.75, which effectively filtered out most river shoreline pixels^39^. After that, non-river-channel and narrow-channel (< ∼60 m) pixels were carefully removed manually, and the SR image was resampled to match the resolution of water occurrence data. Next, the dominant LULC type for each river pixel was determined within a 4-km circle. We computed Chl-a, TSS, and DOC for river pixels following well-established methods^40-42^, and plotted the water property values for dominant LULC types (Fig. 2). We also extracted water property values at eDNA sampling sites and plotted fish species richness against water properties across the catchment (Fig. 5). 14 sites at narrow channels were removed to ensure accuracy of calculations. Detailed formulas of Chl-a, TSS, and DOC calculation can be found in the Supplementary Text.

## Acknowledgements

We especially thank Yixin Hao in space physics for his invaluable insights and inspirations to our FishDiv-LULC modeling framework. We thank Elvira Mächler for her help with eDNA sample processing, Michael O’Brien, Luca Carraro, and Helen Kurkjian for their help with revising the manuscript. F.A. is funded by the Swiss National Science Foundation Grants No. 31003A_173074 and 310030_197410, and F.A, R.F. and M.S. are further supported by the University of Zurich Research Priority Programme on Global Change and Biodiversity (URPP GCB).

## Author Contributions

F.A. and H.Z. designed the research; H.Z. performed the research and developed the model; R.F. contributed to statistical methods; M.O. conducted the fieldwork; R.C.B., M.O., J.B., C.D.M., L.R.H., and B.H. did the bioinformatic analysis; H.Z. and F.A. wrote the paper, and all authors contributed to revising the text.

