## Supplementary Information for "Terrestrial land cover shapes fish diversity in major subtropical rivers"

**Author affiliations**:

**This file includes:**

**Supplementary Text**

**Figures S1 – S11**

**Tables S1 – S9**

**SI References**

### Supplementary Text

#### Calculation of pixel length and area

We adopted the widely-used haversine formula to estimate pixel length ($d_{\mathrm{lon}}^{\mathrm{j}}$, $d_{\mathrm{lat}}^{\mathrm{j}}$) and area ($A_{\mathrm{j}}$) in a map. The formula is specified as follows.

$$a = \sin^2\left(\frac{\varphi_1 - \varphi_2}{2}\right) + \cos\varphi_1 \cos\varphi_2 \sin^2\left(\frac{\lambda_1 - \lambda_2}{2}\right),$$

$$d = R \cdot 2 \cdot \arctan\left(\sqrt{\frac{a}{1-a}}\right),$$

$$A_{\mathrm{j}} = d_{\mathrm{lon}}^{\mathrm{j}} \cdot d_{\mathrm{lat}}^{\mathrm{j}}.$$

Whereby $\lambda_1$, $\lambda_2$ and $\varphi_1$, $\varphi_2$ (in radians) are the longitudes and latitudes of pixel 1 and 2, respectively. *R* is the radius of the Earth, and 6,371 km is used here. Typically, the longitudinal and latitudinal distances ($d_{\mathrm{lon}}$ and $d_{\mathrm{lat}}$) of mesh grid are calculated by choosing pixel 1 and 2 as longitudinal or latitudinal consecutive pixels. Then, $A_{\mathrm{j}}$ is estimated as a product of $d_{\mathrm{lon}}^{\mathrm{j}}$ and $d_{\mathrm{lat}}^{\mathrm{j}}$. The range of pixel area in this catchment is: $8.97 \times 10^{-2} < A_{\mathrm{j}} < 9.27 \times 10^{-2}$ km$^2$.

#### Comparison of distance decay functions

Distance decay is one of the most common phenomena in the natural world, especially in real-world spatial processes. There are multiple widely-used distance decay functions in the field of spatial modeling, including the exponential decay function (exp), the spherical function (sph), the Matérn function (M), and the Gaussian function (G). Here, we compared the performance of these distance decay functions in the FishDiv-LULC modeling. For the Matérn function, we chose smoothness *v* = 1.5 (M15) and 2.5 (M25) as they were “smooth” enough and easy to compute (a product of a polynomial and an exponent) (*1*). These formulas are shown as follows.

$$f_{exp}(d) = \frac{3}{r} e^{-\frac{3d}{r}}.$$

$$f_{sph}(d) = \begin{cases} \frac{3}{8} r \left(1 - \frac{3}{2}\left(\frac{d}{r}\right) + \frac{1}{2}\left(\frac{d}{r}\right)^3\right), & 0 \le d \le r \\ 0, & d > r \end{cases}.$$

$$f_{M15}(d) = \frac{3\sqrt{3}}{2r}\left(1 + \frac{3\sqrt{3}d}{r}\right) e^{-\frac{3\sqrt{3}d}{r}}.$$

$$f_{M25}(d) = \frac{9\sqrt{5}}{8r}\left(1 + \frac{3\sqrt{5}d}{r} + \frac{15d^2}{r^2}\right)e^{-\frac{3\sqrt{5}d}{r}}.$$

$$f_G(d) = 6\sqrt{\frac{1}{\pi r^2}}e^{-\frac{9d^2}{r^2}}.$$

Whereby, *r* is the effective distance; *d* is the flow distance between the pixel and the sampling site. All functions were already normalized to ensure $\int_0^\infty f(x)\,dx = 1$.

We conducted the optimization process by replacing $f(d)$ with the distance decay functions above. The detailed results are shown in Tab. S3. In summary, we found these functions are comparable, therefore we chose the mostly widely-used exponential function in the FishDiv-LULC model.

**Significance and confidence interval of model parameters (likelihood-ratio test)**

We determined the significance of LULC values (***V***), river discharge (coefficient *b* in the model) and effective spatial range (*r*) by conducting a likelihood-ratio test. Specifically, we exclude one parameter at each time, and have hypotheses $H_0$ and $H_1$ as follows.

$$H_0: \boldsymbol{\theta} = \boldsymbol{\theta}_0, \qquad \boldsymbol{\theta}_0 \in \Theta_0.$$

$$H_1: \boldsymbol{\theta} = \boldsymbol{\theta}, \qquad \boldsymbol{\theta} \in \Theta.$$

Whereby, $\Theta_0$ is a subset of parameter space $\Theta$; and in our case, $\Theta \setminus \Theta_0 = \mathcal{V}_i$, $\mathcal{V}_i$ is the space of the excluded parameter. Then, the likelihood ratio test statistic for $H_0$ is given by:

$$\lambda_{LR} = -2\ln\left[\frac{\sup_{\boldsymbol{\theta}\in\Theta_0}\mathcal{L}(\boldsymbol{\theta}|\boldsymbol{B})}{\sup_{\boldsymbol{\theta}\in\Theta}\mathcal{L}(\boldsymbol{\theta}|\boldsymbol{B})}\right] = -2[\ell(\boldsymbol{\mu}(\boldsymbol{\theta}_0),\sigma^2|\boldsymbol{B}) - \ell(\boldsymbol{\mu}(\boldsymbol{\theta}),\sigma^2|\boldsymbol{B})].$$

Under the null hypothesis, $\lambda_{LR}$ converges asymptotically to a $\chi^2$ distribution (Wilks' theorem). The degree of freedom of $\chi^2$ distribution is one in our case as we only exclude one parameter at each time. Finally, we are able to compute the significance level (p-value) based on $\ell(\boldsymbol{\mu}(\boldsymbol{\theta}_0),\sigma^2|\boldsymbol{B})$ and $\ell(\boldsymbol{\mu}(\boldsymbol{\theta}),\sigma^2|\boldsymbol{B})$ and $\chi^2$ distribution.

We further computed 50 % and 90 % profile likelihood-ratio confidence intervals (CI) for each parameter (Table S8) based on likelihood-ratio tests. Briefly, we set values for one parameter, and then compare $\lambda_{LR}$ with a $\chi^2$ distribution (degree of freedom is one) to determine if these values are within a certain level CI.

#### Calculation of river water properties

We employed a cloud-free Sentinel-2 surface reflectance (SR) image and water occurrence data to estimate the water properties of chlorophyll-a content (Chl-a), total suspended solids (TSS), and dissolved organic carbon (DOC) in the major river channels. All methods we applied are well-established on the Sentinel-2 SR data. The detailed calculations and formulas are shown as follows. Here, BX represents the corresponding band of Sentinel-2 SR image.

##### Chlorophyll-a content (Chl-a)

Chl-a content is a direct indicator for photosynthesis activities in river, thus an accurate proxy for productivity in aquatic ecosystems. We adopted a blending method ($Chl_{\mathrm{Blend}}$) combining OCx and 2-Band algorithms to calculate Chl-a (mg/m$^3$), because of its good accuracy and worldwide adaptability (*2-4*).

$$x = \log_{10}(\max(\mathrm{B1}, \mathrm{B2})/\mathrm{B3})$$

$$y = 0.3308 - 2.6684x + 1.5990x^2 + 0.5525x^3 - 1.4876x^4$$

$$\boldsymbol{Chl_{\mathrm{OC3}} = 10^{y}}.$$

$$\mathrm{CI} = \mathrm{B3} - 0.473 \cdot \mathrm{B1} - 0.527 \cdot \mathrm{B4}$$

$$\boldsymbol{Chl_{\mathrm{CI}} = 10^{-0.4909+191.6590 \cdot \mathrm{CI}}}.$$

$$b = \max(0.15, \min(Chl_{\mathrm{CI}}, 0.20))$$

$$w1 = (b - 0.15)/(0.20 - 0.15)$$

$$\boldsymbol{Chl_{\mathrm{OCx}} = w1 \cdot Chl_{\mathrm{OC3}} + (1 - w1) \cdot Chl_{\mathrm{CI}}}.$$

$$\boldsymbol{Chl_{\mathrm{2-Band}} = (35.75 \cdot \mathrm{B5}/\mathrm{B4} - 19.3)^{1.124}}.$$

$$r = \max(0.75, \min(\mathrm{B5}/\mathrm{B4}, 1.15))$$

$$w2 = (r - 0.75)/(1.15 - 0.75)$$

$$\boldsymbol{Chl_{\mathrm{Blend}} = w2 \cdot Chl_{\mathrm{2-Band}} + (1 - w2) \cdot Chl_{\mathrm{OCx}}}.$$

##### Total suspended solids (TSS)

TSS are particles in water larger than 2 microns, and are related to turbidity of the river. They come from soil, sediments, decaying organic matter, wastewater, etc., indicating erosion, pollution, and availability of organic matters. We used a method proposed by (*5*) to estimate TSS (g/m$^3$) with Sentinel-2 SR Band 4.

$$TSS = 1.74 + (355.85 \cdot \pi \cdot \mathrm{B4})/(1 - \pi \cdot \mathrm{B4}/1728).$$

**Dissolved organic carbon (DOC)**

DOC, coming from algae, soil, aquatic or terrestrial plants or animals, is a basic form of nutrient supporting the growth of organisms in river, often related to the amount of organic matters or materials in aquatic systems. We applied a method by (*6*) to estimate DOC concentration in river. This method first retrieves the colored dissolved organic matter (CDOM) absorption coefficient at 440 nm, namely $a_{\mathrm{CDOM}}(440)$, then calculated DOC (mg/L) using the relationship between $a_{\mathrm{CDOM}}(440)$ and DOC.

$$a_{\mathrm{CDOM}}(440) = 28.966 \cdot e^{-2.015 \cdot \mathrm{B3}/\mathrm{B4}}.$$

$$DOC = 1.268 \cdot a_{\mathrm{CDOM}}(440) + 3.623.$$

#### Figures

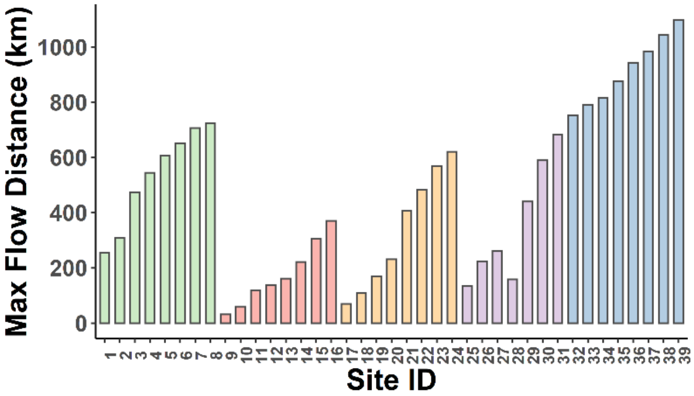


**Figure S1** Maximum flow distance for sampling sites in the Chao Phraya catchment. Flow distance was calculated from the flow direction map of HydroSHEDS (version 1). Colors indicate major tributaries in this catchment (Ping River in green, Wang River in red, Yom River in yellow, Nan River in purple, and lower reaches of Chao Phraya River in blue).

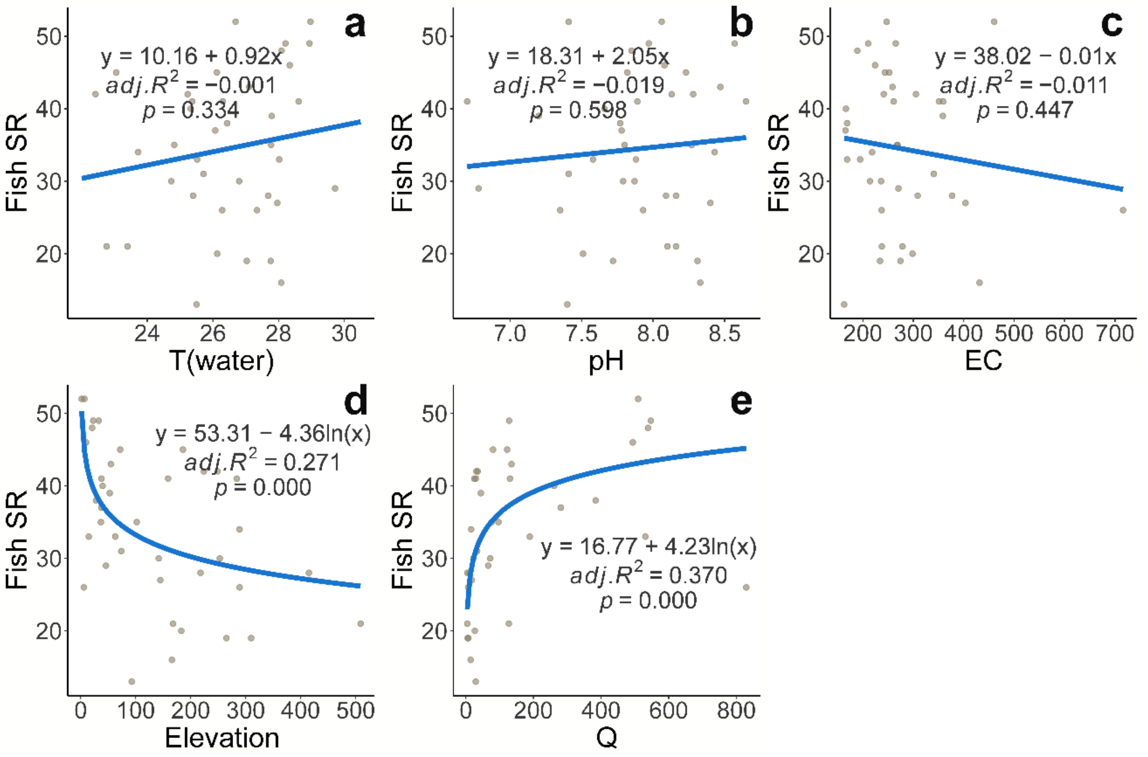


**Figure S2** Scatter plots of fish species richness against environmental variables, including water temperature (**a**), pH (**b**), electricity conductivity (**c**), elevation (ln-transformed) (**d**), and river discharge (ln-transformed) (**e**). Results of regression analyses are shown. Among all the five variables, flow discharge explains fish species richness the most in the catchment.

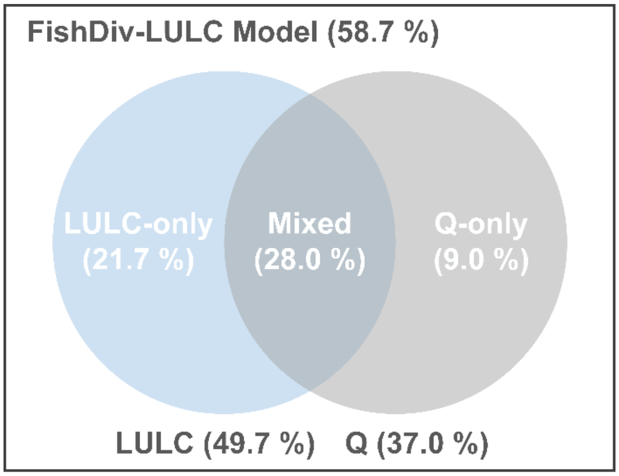


**Figure S3** Variance partitioning of the FishDiv-LULC model in the Chao Phraya catchment. We fitted a LULC model (excl. discharge) and a discharge (Q) model (excl. LULC), and then compared them with FishDiv-LULC modeling results. Values in the figure are adjusted-$R^2$.

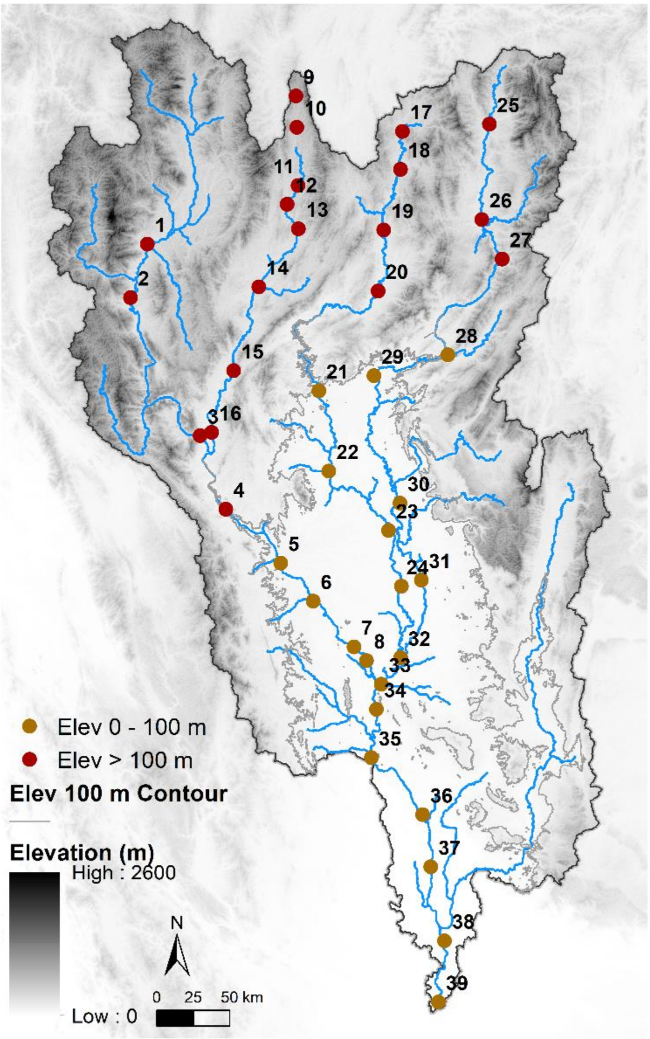


**Figure S4** Division of mountainous and plain sites. The elevation of sampling sites ranges from 2 to 509 m a.s.l. A contour of 100 m elevation is shown (encircling all areas lower than 100 m as a white polygon) is used as the criterion of separation. 19 sites fall in the "mountainous" region (colored in red), while 20 sites in the plain region (colored in dark yellow).

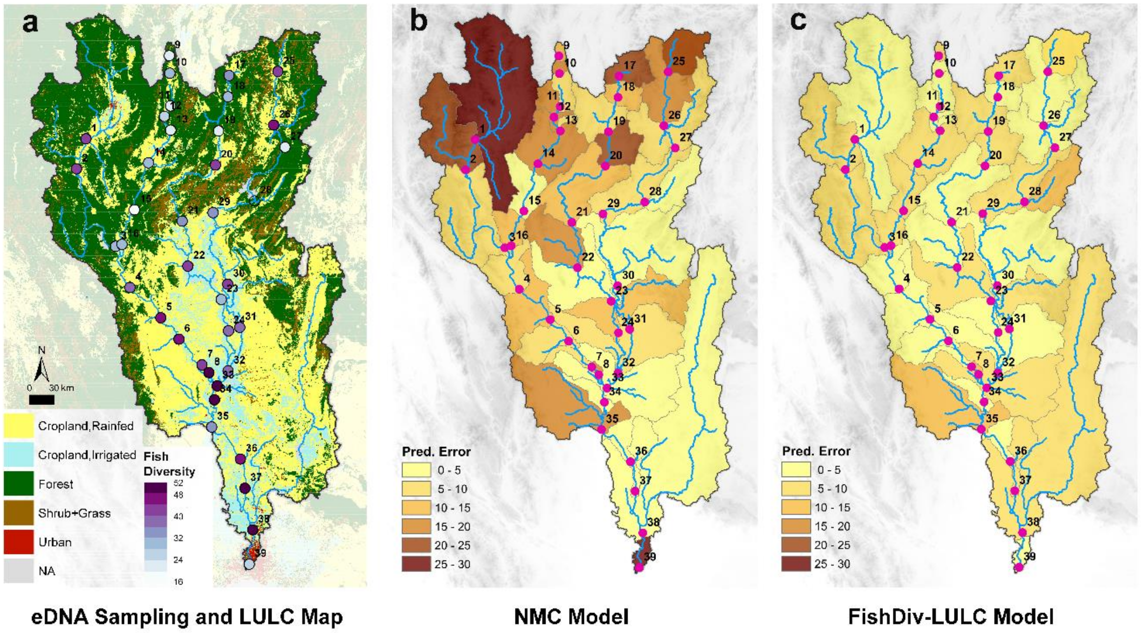


**Figure S5** Comparison of prediction errors between a neutral meta-community (NMC) model (**b**) and the FishDiv-LULC model (**c**). In the case of Chao Phraya catchment, the FishDiv-LULC model better captures fish species richness pattern than NMC model, especially in the hilly and mountainous region. The optimal parameter values for NMC model are shown in Tab. S3.

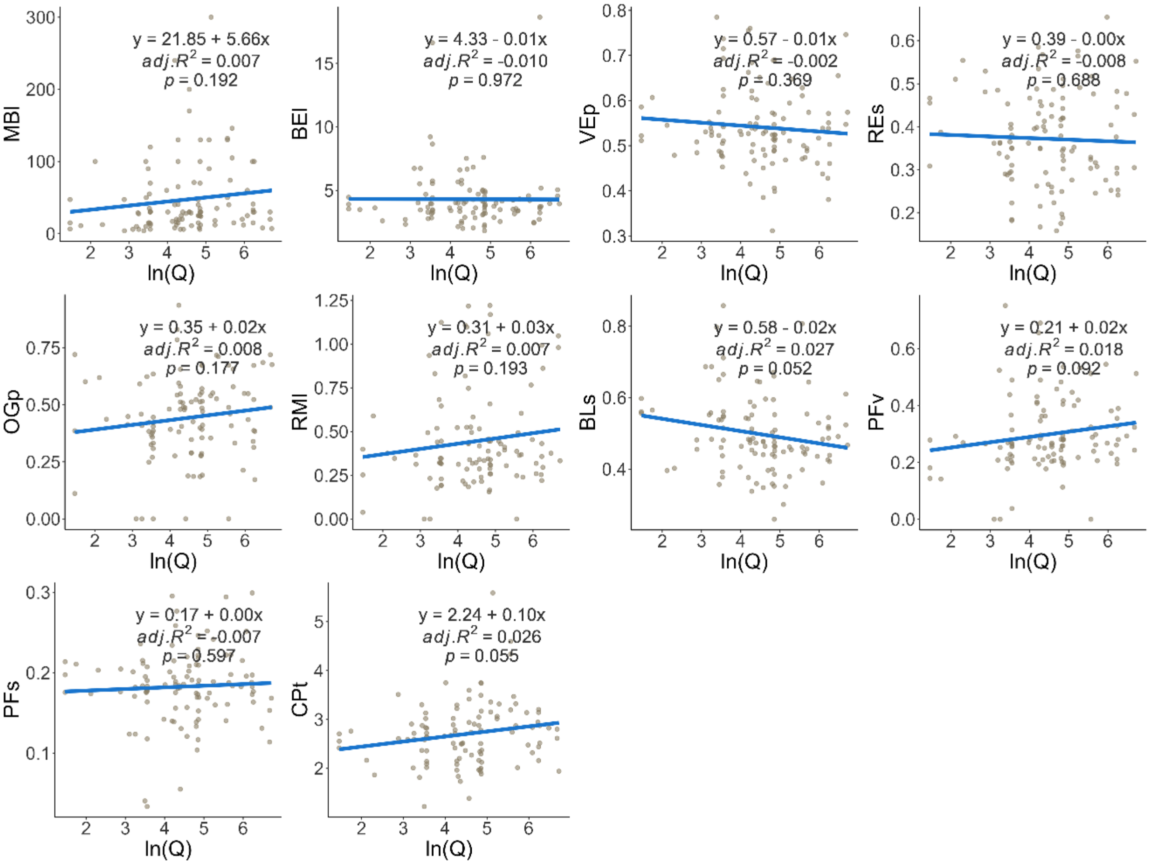


**Figure S6** Scatter plots of fish traits against river discharge, a proxy of river channel characteristics. Ten major fish morphological traits were collected from the FISHMORPH database: maximum body length (MBl), body elongation (BEl), vertical eye position (VEp), relative eye size (REs), oral gape position (OGp), relative maxillary length (RMl), body lateral shape (BLs), pectoral fin vertical position (PFv), pectoral fin size (PFs), and caudal peduncle throttling (CPt). River discharge for each species was derived by the median of all discharge values at species occurrence sites.

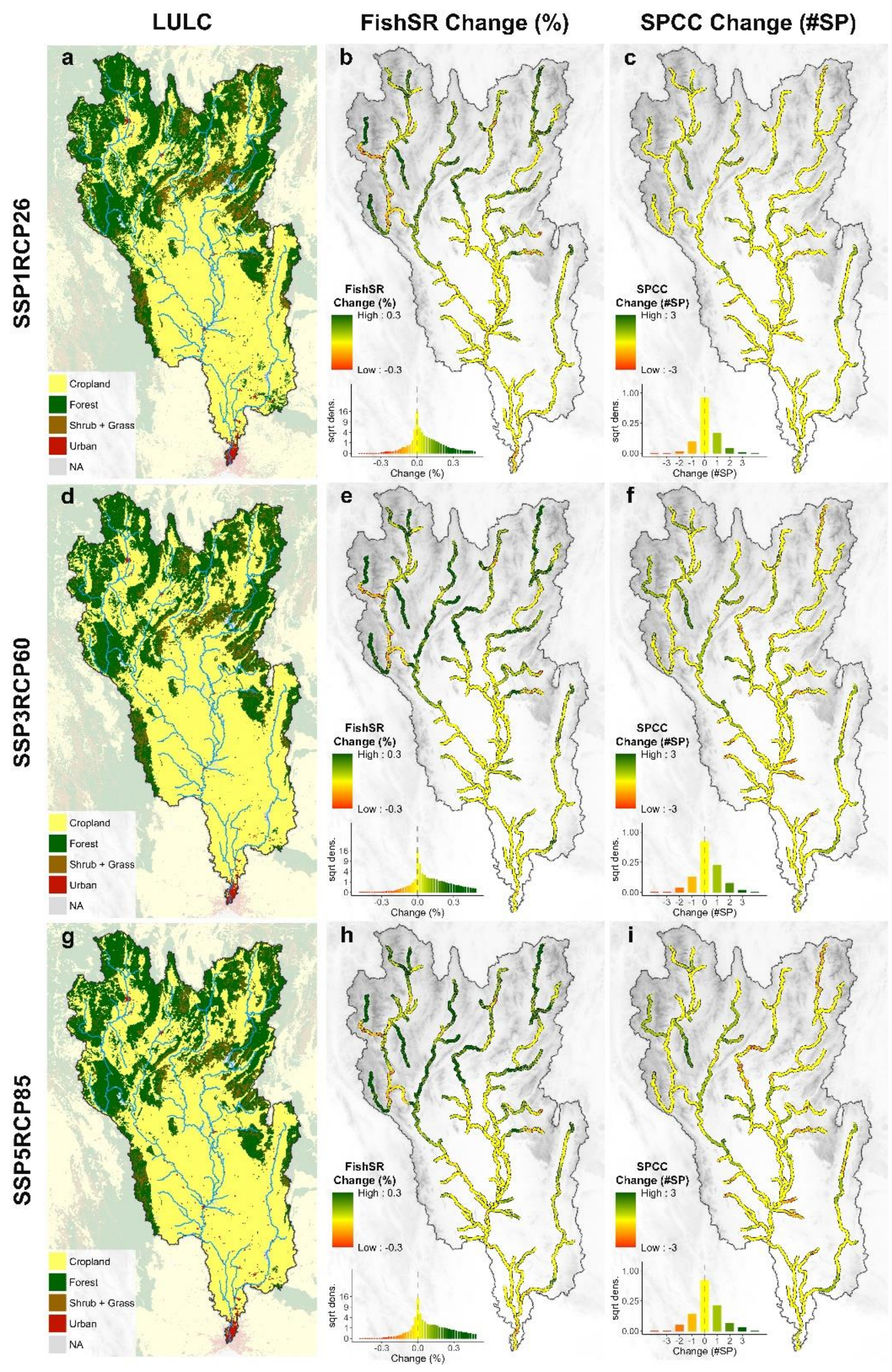


**Figure S7** GLOBIO4 predicted LULC maps in 2050 under SSP1RCP26 (**a**), SSP3RCP60 (**d**), and SSP5RCP85 (**g**) scenarios. We projected percentages of fish species richness change (**b**, **e**, and **h**) and changes of fish species of conservational concern (SPCC) (**c**, **f**, and **i**) in 2016—2050. We observed an overall rise of fish species richness in the mountainous area, but divergent patterns in the change of SPCC.

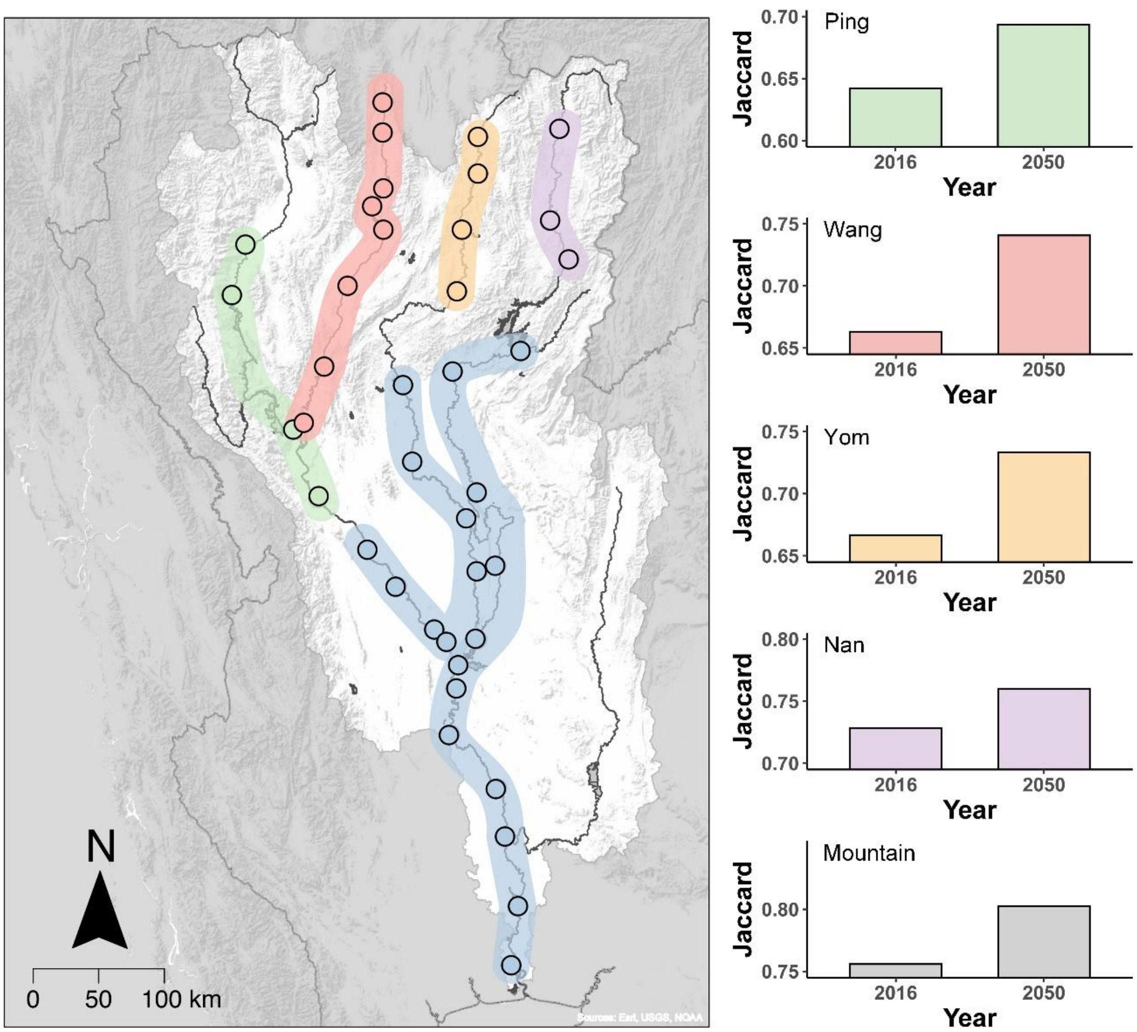


**Figure S8** Jaccard similarity values of fish assemblages between sites in upper reaches of Ping, Wang, Yom, and Nan (elevation ≥ 100 m) and the plain (elevation < 100 m). We compared fish assemblages of 2016 and 2050 under SSP5RCP85 according to our model predictions, and found a notable increase of Jaccard similarity in the future. These results illustrate homogenization of fish assemblages after cropland expansion in the mountains in 2050.

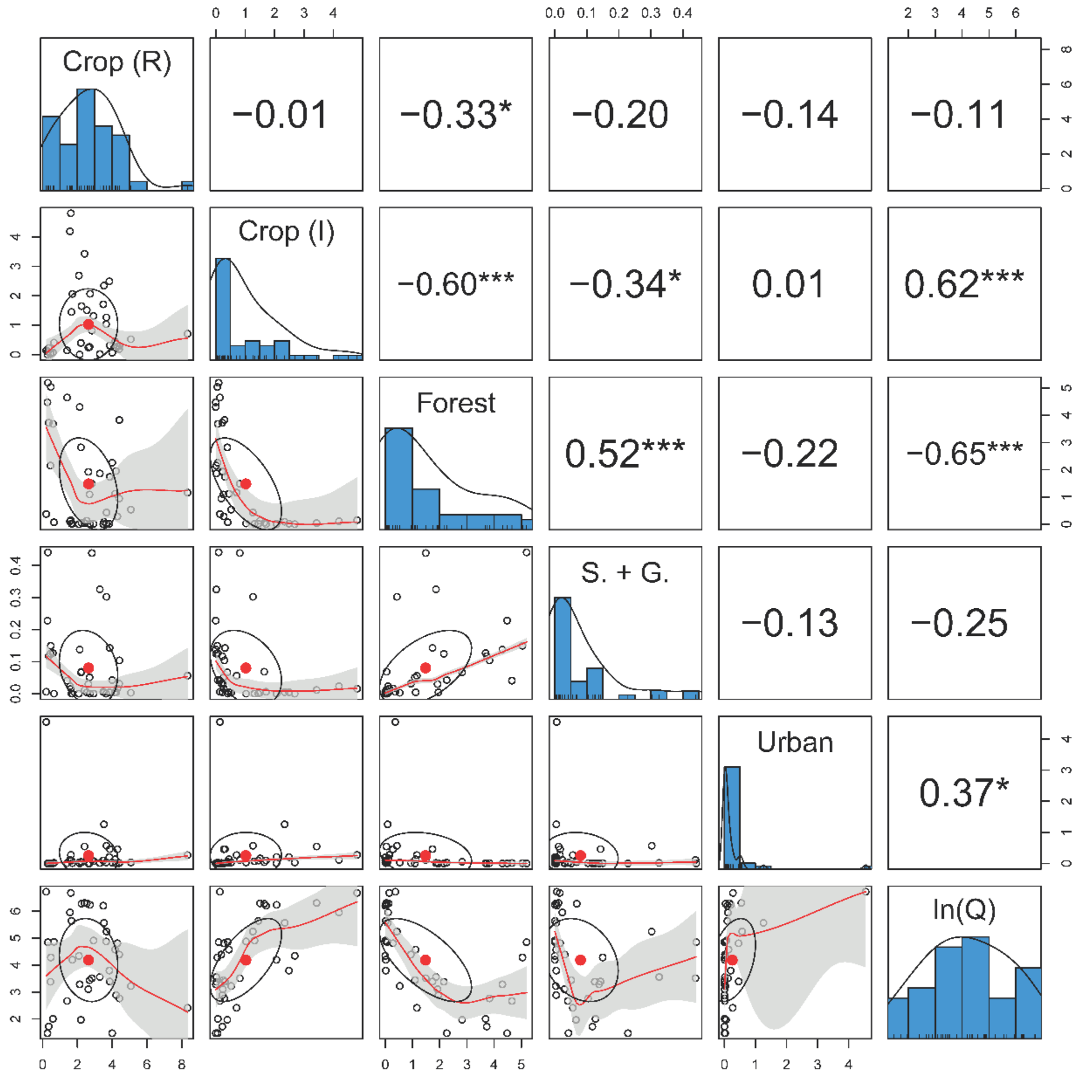


**Figure S9** Paired correlation plot of LULC types and river discharge across Chao Phraya catchment. The plot was calculated based on the catchment-dependent ***C***(*r*) matrix with the optimal effective distance *r* = 19 km. The results show no high correlation among the predictors.

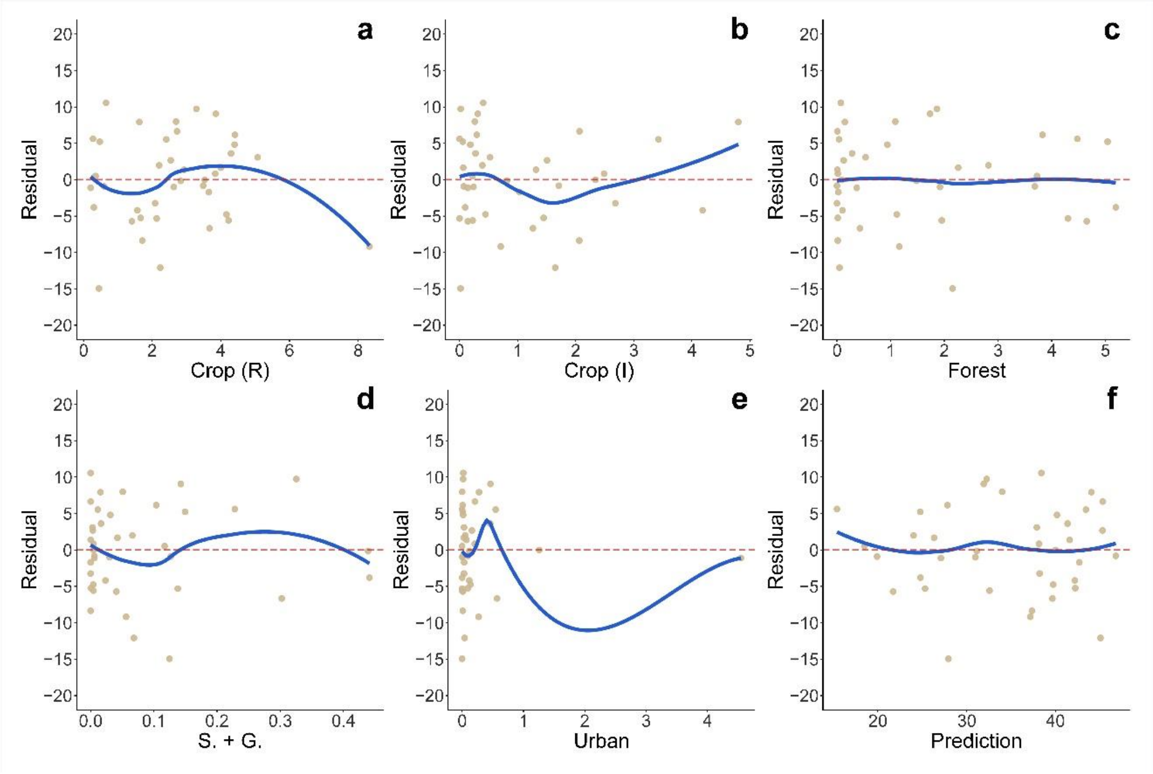


179 **Figure S10** Residuals against LULC effects (**a**—**e**) and model predictions (**f**). LULC effect values for
180 each site of (**a**) rainfed cropland, (**b**) irrigated cropland, (**c**) forest, (**d**) shrub- and grassland, and
181 (**e**) urban areas were calculated based on the catchment-dependent ***C***(*r*) matrix with the optimal
182 effective distance *r* = 19 km. No obvious trend was found in these scatter plots. Therefore, the
183 FishDiv-LULC model is generally unbiased.

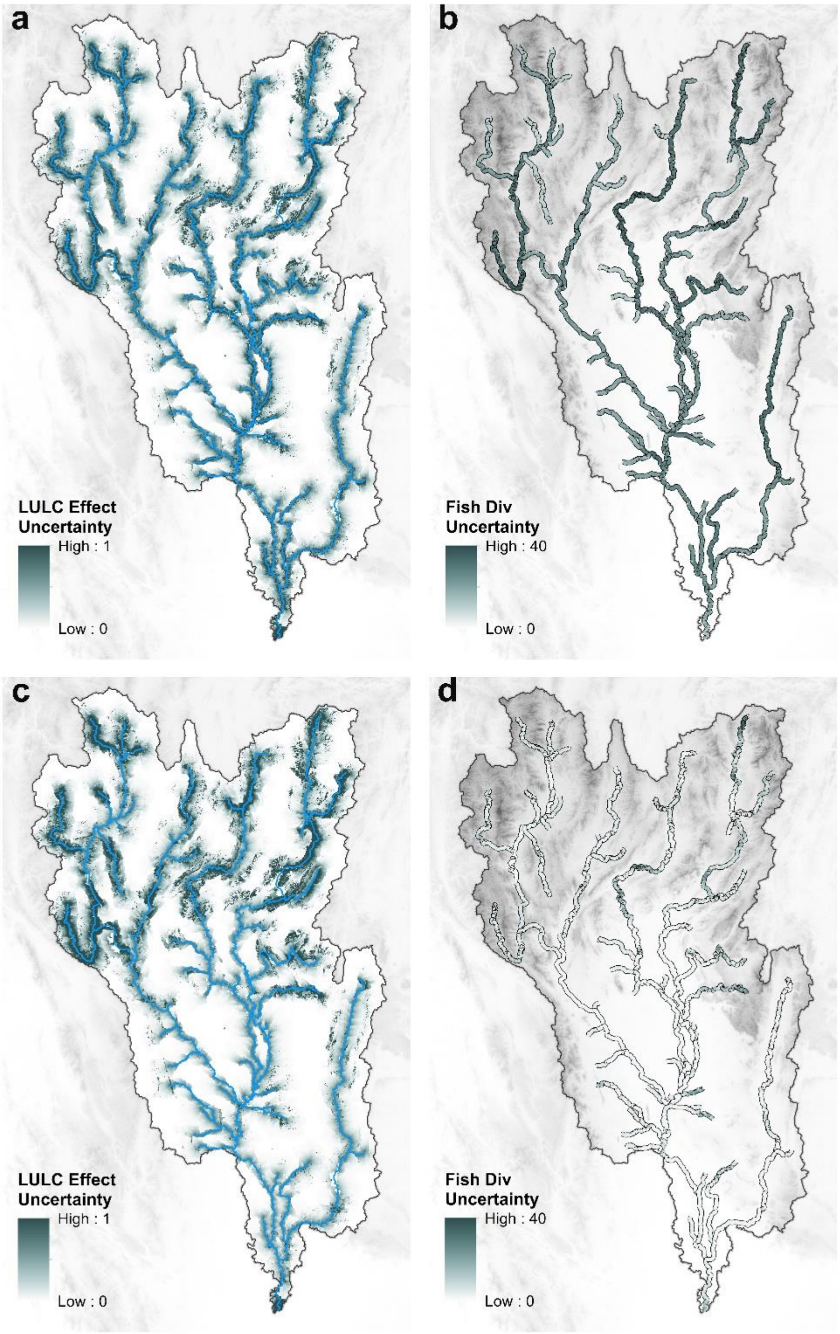


**Figure S11** Uncertainty maps of LULC effects and river fish diversity predictions. (**a**) and (**b**) were calculated based on five-class LULC system; while (**c**) and (**d**) were uncertainties of four-class LULC system, in which rainfed cropland and irrigated cropland were merged. The uncertainty was estimated as the interquartile range (IQR) of predictions from 2,000 bootstrapped models.

### Tables

**Table S1** Recoding table of land use and land cover (LULC) types of the European Space Agency Climate Change Initiative (ESA CCI) land cover map. We merged the original land cover map into a five-class-system map according to this table.

| ESA CCI Value | ESA CCI Label | Area (%) | Five-Class System |
|---|---|---|---|
| 10 | Cropland, rainfed | 35.91% | Cropland, Rainfed |
| 11 | Cropland, rainfed, herbaceous cover | 3.58% | Cropland, Rainfed |
| 12 | Cropland, rainfed, tree or shrub cover | < 0.01% | Cropland, Rainfed |
| 20 | Cropland, irrigated or post-flooding | 9.55% | Cropland, Irrigated |
| 30 | Mosaic cropland (>50%) / natural vegetation (tree, shrub, herbaceous cover) (<50%) | 3.15% | Cropland, Rainfed |
| 40 | Mosaic natural vegetation (tree, shrub, herbaceous cover) (>50%) / cropland (<50%) | 5.07% | Forest |
| 50 | Tree cover, broadleaved, evergreen, closed to open (>15%) | 9.72% | Forest |
| 60 | Tree cover, broadleaved, deciduous, closed to open (>15%) | 5.73% | Forest |
| 61 | Tree cover, broadleaved, deciduous, closed (>40%) | 0.02% | Forest |
| 62 | Tree cover, broadleaved, deciduous, open (15-40%) | — | — |
| 70 | Tree cover, needle-leaved, evergreen, closed to open (>15%) | 2.97% | Forest |
| 71 | Tree cover, needle-leaved, evergreen, closed (>40%) | — | — |
| 72 | Tree cover, needle-leaved, evergreen, open (15-40%) | — | — |
| 80 | Tree cover, needle-leaved, deciduous, closed to open (>15%) | < 0.01% | Forest |
| 81 | Tree cover, needle-leaved, deciduous, closed (>40%) | — | — |
| 82 | Tree cover, needle-leaved, deciduous, open (15-40%) | — | — |
| 90 | Tree cover, mixed leaf type (broadleaved and needle-leaved) | — | — |
| 100 | Mosaic tree and shrub (>50%) / herbaceous cover (<50%) | 16.31% | Forest |
| 110 | Mosaic herbaceous cover (>50%) / tree and shrub (<50%) | < 0.01% | Shrub + Grass |
| 120 | Shrubland | 1.12% | Shrub + Grass |
| 121 | Evergreen shrubland | 5.98% | Shrub + Grass |
| 122 | Deciduous shrubland | < 0.01% | Shrub + Grass |
| 130 | Grassland | 0.21% | Shrub + Grass |
| 140 | Lichens and mosses | — | — |
| 150 | Sparse vegetation (tree, shrub, herbaceous cover) (<15%) | — | — |
| 152 | Sparse shrub (<15%) | — | — |
| 153 | Sparse herbaceous cover (<15%) | — | — |
| 160 | Tree cover, flooded, fresh or brackish water | — | — |
| 170 | Tree cover, flooded, saline water | < 0.01% | Forest |
| 180 | Shrub or herbaceous cover, flooded, fresh/saline/brackish water | 0.03% | Shrub + Grass |
| 190 | Urban areas | 0.65% | Urban |
| 200 | Bare areas | < 0.01% | — |
| 201 | Consolidated bare areas | — | — |
| 202 | Unconsolidated bare areas | — | — |
| 210 | Water bodies | < 0.01% | — |
| 220 | Permanent snow and ice | — | — |

**Table S2** The estimated parameter values of separate mountainous and plain area models. The 39 sampling sites were classified as mountainous (19 sites) or plain (20 sites) by if the elevation of sites is greater than 100 m. P values were computed by the likelihood-ration test which eliminated a single predictor at each time.

| | a | Crop (R) | Crop (I) | Forest | S. + G. | Urban | b | *r* (km) | adj.$R^2$ | $-2\ell$ |
|---|---|---|---|---|---|---|---|---|---|---|
| **Mountain** | 30.618 | 1.000 | -2.248 | -3.230 | 12.033 | -11.440 | 2.779 | 32 | 0.614 | 109.840 |
| **p value** | — | 0.293 | 0.833 | 0.001 | 0.209 | 0.633 | 0.014 | <0.001 | — | — |
| **Plain** | 14.150 | 3.656 | -0.677 | -5.038 | -173.65 | -4.828 | 4.690 | 10 | 0.562 | 121.338 |
| **p value** | — | 0.012 | 0.629 | 0.253 | 0.027 | 0.002 | 0.003 | <0.001 | — | — |
| **Whole** | 20.585 | 1.438 | -0.238 | -2.163 | -4.857 | -3.684 | 3.550 | 19 | 0.587 | 250.613 |
| **p value** | — | 0.054 | 0.852 | 0.055 | 0.631 | 0.019 | 0.003 | <0.001 | — | — |

**Table S3** The optimal parameters for NMC models in the case of Chao Phraya catchment. Parameters a and b control the dispersion of fish unit. $w_u$ is the weight factor modifying the upstream distance. $C_H$ is the average habitat capacity in a direct tributary area. *v* is the fish species diversification rate. For details of the NMC model, please refer to (*7*).

| **a** | **b** | $w_u$ | $C_H$ | ***v*** | adj.$R^2$ |
|---|---|---|---|---|---|
| 0.832 | 0.125 | 1.828 | 48 | 1.07e-5 | 0.255 |

**Table S4** Mean (**a**) and range (**b**) of fish traits in relation to LULC. We collected all major fish morphological traits related to functioning from FISHMORPH database. The traits are maximum body length (MBl), body elongation (BEl), vertical eye position (VEp), relative eye size (REs), oral gape position (OGp), relative maxillary length (RMl), body lateral shape (BLs), pectoral fin vertical position (PFv), pectoral fin size (PFs), and caudal peduncle throttling (CPt). The area of 2D fish trait envelopes is shown in (**c**).

(**a**) Mean

| LULC | MBl | BEl | VEp | REs | OGp | RMl | BLs | PFv | PFs | CPt |
|---|---|---|---|---|---|---|---|---|---|---|
| **Crop (R)** | 39.5 | 4.54 | 0.54 | 0.37 | 0.42 | 0.43 | 0.49 | 0.28 | 0.18 | 2.58 |
| **Crop (I)** | 58.5 | 3.88 | 0.56 | 0.38 | 0.53 | 0.58 | 0.48 | 0.34 | 0.18 | 2.84 |
| **Forest** | 24.7 | 4.10 | 0.59 | 0.33 | 0.42 | 0.46 | 0.60 | 0.29 | 0.20 | 2.29 |
| **S. + G.** | 34.5 | 3.98 | 0.53 | 0.41 | 0.43 | 0.36 | 0.49 | 0.28 | 0.19 | 2.81 |
| **Urban** | 87.5 | 5.11 | 0.49 | 0.34 | 0.41 | 0.34 | 0.47 | 0.33 | 0.17 | 2.95 |

(**b**) Range

| LULC | MBl | BEl | VEp | REs | OGp | RMl | BLs | PFv | PFs | CPt |
|---|---|---|---|---|---|---|---|---|---|---|
| **Crop (R)** | 126 | 16.6 | 0.36 | 0.37 | 0.72 | 1.17 | 0.51 | 0.54 | 0.22 | 2.53 |
| **Crop (I)** | 234 | 4.47 | 0.35 | 0.44 | 0.76 | 1.07 | 0.35 | 0.49 | 0.19 | 2.51 |
| **Forest** | 96 | 5.60 | 0.30 | 0.25 | 0.37 | 0.66 | 0.33 | 0.64 | 0.17 | 1.49 |
| **S. + G.** | 140 | 4.98 | 0.26 | 0.36 | 0.78 | 0.77 | 0.33 | 0.55 | 0.24 | 1.87 |
| **Urban** | 293 | 14.1 | 0.40 | 0.38 | 0.68 | 0.52 | 0.51 | 0.51 | 0.26 | 4.20 |

(**c**) Area of 2D envelopes

| Crop (R) | Crop (I) | Forest | S. + G. | Urban |
|---|---|---|---|---|
| 21.4 | 16.28 | 15.4 | 16.1 | 24.5 |

**Table S5** List of fish species of conservation concern (SPCC) for which we detected an eDNA signal in the Chao Phraya River catchment. The endangered level is obtained from the red list of the International Union for Conservation of Nature (IUCN). CR, EN, VU, and NT represent critically endangered, endangered, vulnerable, and near threatened, respectively. *Cyprinus carpio* (Eurasian carp) and *Hypophthalmichthys molitrix* (silver carp) were removed because they are alien or invasive species in this catchment.

| SPCC ID | Species Name | Endangered Level |
|---|---|---|
| 1 | *Pangasianodon gigas* | CR |
| 2 | *Pangasianodon hypophthalmus* | EN |
| 3 | *Wallago attu* | VU |
| 4 | *Epalzeorhynchos bicolor* | CR |
| 5 | *Yasuhikotakia sidthimunki* | EN |
| 6 | *Cirrhinus molitorella* | NT |
| 7 | *Rhinogobius chiengmaiensis* | VU |

**Table S6** The comparison of performance of five distance decay functions in the FishDiv-LULC model. The distance decay functions include the exponential function (exp), the spherical function (sph), the Matérn functions with smoothness $v$ = 1.5 (M15) and 2.5 (M25) and the Gaussian function (G). There is no statistical difference between models.

| | a | Crop (R) | Crop (I) | Forest | S. + G. | Urban | b | *r* (km) | *adj.R*$^2$ | $-2\ell$ |
|---|---|---|---|---|---|---|---|---|---|---|
| **exp** | 20.585 | 1.438 | -0.238 | -2.163 | -4.857 | -3.684 | 3.550 | 19 | 0.587 | 250.613 |
| **sph** | 21.695 | 1.026 | -0.088 | -1.623 | -4.088 | -3.272 | 3.338 | 31 | 0.593 | 250.020 |
| **M15** | 21.011 | 1.233 | -0.195 | -1.903 | -4.138 | -3.396 | 3.474 | 25 | 0.589 | 250.367 |
| **M25** | 21.011 | 1.274 | -0.201 | -1.966 | -4.273 | -3.507 | 3.474 | 25 | 0.589 | 250.367 |
| **G** | 21.473 | 1.057 | -0.188 | -1.756 | -3.750 | -3.233 | 3.426 | 24 | 0.592 | 250.132 |

**Table S7** Variance inflation factors (VIFs) of LULC types and flow discharge across Chao Phraya catchment. VIFs were calculated based on the catchment-dependent $\boldsymbol{C}(r)$ matrix under the optimal effective distance *r* = 19 km. The results show no high correlation among predictive parameters as all VIF < 5.

| | Crop (R) | Crop (I) | Forest | S. + G. | Urban | ln(Q) |
|---|---|---|---|---|---|---|
| **VIF** | 1.472 | 2.111 | 3.279 | 1.422 | 1.328 | 2.752 |

**Table S8** Profile likelihood ratio confidence intervals (CIs) of levels of 50 % and 90 % of FishDiv-LULC model parameters.

| | Crop (R) | Crop (I) | Forest | S. + G. | Urban | b | *r* (km) |
|---|---|---|---|---|---|---|---|
| **5%** | 0.245 | -2.622 | -4.465 | -40.337 | -6.462 | 1.640 | 11 |
| **25%** | 0.945 | -1.069 | -2.947 | -13.329 | -4.737 | 2.747 | 16 |
| **value** | 1.438 | -0.238 | -2.163 | -4.857 | -3.684 | 3.550 | 19 |
| **75%** | 1.976 | 0.558 | -1.423 | 1.879 | -2.612 | 4.300 | 24 |
| **95%** | 3.130 | 1.775 | -0.387 | 15.311 | -1.124 | 5.478 | 34 |

**Table S9** Estimations of the four-LULC-class FishDiv-LULC model parameters with the spatial range *r*. Cropland (rainfed) and cropland (irrigated) were merged in the four-class LULC system. The magnitude of LULC effect of cropland, forest, shrub + grass, and urban areas are provided. $\ell$ is the log-likelihood function for optimization, with $-2\ell$ increasing by 1.7 compared to the five-class model. The significance of parameters is determined by a likelihood-ratio test.

| | **a** | **Crop.** | **Forest** | **S. + G.** | **Urban** | **b** | ***r* (km)** | **adj.$R^2$** | $-2\ell$ |
|---|---|---|---|---|---|---|---|---|---|
| **Value** | 23.332 | 1.038 | -2.098 | -2.999 | -3.178 | 2.756 | 20 | 0.582 | 252.320 |
| **p** | — | 0.123 | 0.050 | 0.739 | 0.037 | 0.007 | <0.001 | — | — |
